# Dietary Protection Against the Visual and Motor Deficits Induced by Experimental Autoimmune Encephalomyelitis

**DOI:** 10.1101/2022.07.13.499958

**Authors:** Katarzyna Zyla-Jackson, Dorothy A. Walton, Kendra S. Plafker, Susan Kovats, Constantin Georgescu, Richard S. Brush, Madison Tytanic, Martin-Paul Agbaga, Scott M. Plafker

## Abstract

Five to eight percent of the world population currently suffers from at least one autoimmune disorder. Despite multiple immune modulatory therapies for autoimmune demyelinating diseases of the central nervous system, these treatments can be limiting for subsets of patients due to adverse effects and expense. To circumvent these barriers, we investigated a nutritional intervention in mice undergoing experimental autoimmune encephalomyelitis (EAE). This model of autoimmune-mediated demyelination induces visual and motor pathologies similar to those experienced by people with multiple sclerosis (MS). Here, we report that limiting dietary carbohydrates by feeding mice a ketogenic diet (KD) enriched in medium chain triglycerides, α-linolenic acid, and fiber is effective at mitigating EAE-induced optic neuritis and motor deficits. The KD was efficacious when fed as a preventive regimen prior to EAE immunization as well as when initiated as an interventional regimen following the onset of EAE symptoms. The KD minimally impacted body weight during the experimental time course, increased circulating ketones, prevented motor and ocular deficits, preserved myelination of the optic nerve, and reduced infiltration of immune cells to optic nerves. The KD also increased anti-inflammatory-associated omega-3 fatty acids in the plasma and reduced select cytokines in the circulation associated with EAE-mediated pathological inflammation. In light of ongoing clinical trials using dietary strategies to treat people with MS, these findings support that a KD promotes a systemic anti-inflammatory milieu and ameliorates autoimmune-induced demyelinating visual and motor deficits.

## INTRODUCTION

Multiple sclerosis (MS) is an autoimmune demyelinating disease of the central nervous system (CNS) that causes severe disabilities. Deficits include losses of mobility, balance and coordination, blindness, depression, fatigue, memory loss, and decreased quality of life. MS is more common in females, with onset typically occurring in the third and fourth decades of life (reviewed in (1–3)).

The increased incidence of MS and other autoimmune disorders in recent decades coincides with global increases in obesity, hyperglycemia, hyperinsulinemia, insulin resistance, dyslipidemia, and type 2 diabetes (4). Mounting evidence points to the excessive consumption of ultra-processed foods containing highly-processed carbohydrates and pro-inflammatory fats as drivers of metabolic syndrome (5–7), poor health outcomes (8, 9), and all-cause mortality (10). These ultra-processed, hyperpalatable, calorically-dense foodstuffs promote hyperphagia (11), and can exacerbate autoimmunity by disrupting microbiome-host symbiosis and promoting systemic inflammation (12). Evidence from clinical trials and animal studies support that diet profoundly impacts MS severity and disease trajectory (13–15), consistent with the observation that insulin resistance and adiposity correlate with more severe disability scores for people with MS (16, 17).

To determine the impact of reducing carbohydrate-laden foods in the diet, we investigated the therapeutic efficacy of a ketogenic diet (KD) in the mouse MOG_35-55_-experimental autoimmune encephalomyelitis (MOG_35-55_-EAE) model, hereafter referred to as ‘EAE’. This model of autoimmune-mediated demyelination induces visual and motor pathologies similar to those experienced by people with MS. Despite the caveats and limitations of pre-clinical models (e.g., the lack of genetic diversity inherent to humans), EAE rodent studies have proven valuable with respect to the development of FDA-approved therapeutics for MS (18).

The studies presented here were done to determine whether the KD developed by D’Agostino and colleagues (19) that is enriched in fiber and contains medium chain triglycerides (MCTs; caprylic acid (C8) and capric acid (C10)) along with flaxseed oil and canola oil as the sources of fat can preserve motor and visual function in both male and female C57BL/6J mice undergoing EAE. Previous studies of nutritional interventions using this EAE model have demonstrated that caloric restriction attenuates EAE motor deficits and correlates with reduced levels of IL-6 and leptin (20). Likewise, fast-mimicking diets (21, 22), intermittent fasting (23, 24), and KDs (22, 25, 26) reduce EAE motor disabilities, and a KD improved long-term potentiation, spatial learning, and memory (26).

Nutritional ketosis has been used since the 1920s to treat children with drug-refractory seizures (27, 28), and KDs have gained popularity for weight loss and as an anti-diabetic strategy because the diet suppresses hunger and reduces energy intake (29–31). These and other findings (reviewed in (32, 33)) have led to multiple clinical trials (e.g., NCT01538355, NCT03718247, NCT01915433, NCT05007483) including “Nutritional Approaches in Multiple Sclerosis” (NAMS; NCT03508414)), a randomized controlled clinical trial in Germany for people with active MS to compare a KD versus a fasting protocol versus a fat-modified standard diet over an 18-month period (34).

KDs can differ in composition but share at least two properties. The diets are comprised primarily of fats with moderate protein content and low amounts of carbohydrates, typically in the range of 0-70 g daily for humans, excluding indigestible fiber. Secondly, KDs induce the liver to produce the ketone bodies β-hydroxybutyrate, acetoacetate, and acetone. People in nutritional ketosis have circulating ketones of 0.5 - 4mM with blood glucose less than 150mg/dL, irrespective of fasting or fed state. This nutritional ketosis is distinguished from the pathological ketoacidosis associated with uncontrolled diabetes that yields ketones exceeding 25mM and blood glucose greater than 240mg/dL.

Here we report that a KD can robustly protect against EAE-mediated motor and vision loss concomitant with reducing immune cell infiltration and preserving myelination of the optic nerve in both female and male mice. Functional preservation and protection against neurodegeneration were robust when the KD was fed as a preventive regimen prior to immunization to induce EAE and likewise when implemented as an intervention after symptom onset, demonstrating the translational feasibility of this nutritional approach. The KD increased circulating levels of multiple omega 3 (ω3) fatty acids associated with endogenous resolution pathways of acute inflammation and reduced circulating factors associated with neutrophil-mediated inflammation and MS pathogenicity. Together, these findings show that an MCT-based KD enriched in fiber confers neuroprotection and can reverse the loss of motor and visual function caused by autoimmune-mediated demyelination.

## MATERIALS AND METHODS

### Mice

Male and female C57BL/6J mice were housed in microisolator cages (n=4-5 per cage) under a 12-hr light/dark cycle and fed *ad libitum* (Picolab^®^ Rodent Diet (cat # 5053)). All animal care and experimental procedures were performed in compliance with ARRIVE guidelines, an Oklahoma Medical Research Foundation Institutional Animal Care and Use Committee (IACUC)-approved protocol, and complied with standards delineated by the Institute for Laboratory Animal Research. These studies adhered to The Association for Research in Vision and Ophthalmology (ARVO) statement for the Use of Animals in Research. All studies used C57BL/6J female and male mice. Breeders were purchased from Jackson Laboratories (stock # 000664). At the termination of experiments, mice were anesthetized to collect blood by cardiac puncture and subsequently sacrificed by CO2 asphyxiation followed by cervical dislocation.

### Diet compositions

Teklad control (TD.170645; CD) and ketogenic (TD.10911; KD) diets were obtained from Envigo, Inc., and custom formulated with the assistance of a company nutritionist based on (19). At the macronutrient level, the KD provides 4.7 Kcal/g with 22.4% Kcal from protein, 0.5% Kcal from carbohydrate, and 77.1% Kcal from fat. The CD provides 3.6 Kcal/g with 20.4% Kcal from protein, 69.3% Kcal from carbohydrate, and 10.4% Kcal from fat. Macromolecular compositions and ingredients for both diets are provided in Fig. 1A and Supplemental Fig. 1A. Food and water were provided *ad libitum* and additional hydration/electrolytes were provided to animals showing signs of dehydration using intraperitoneal saline injections.

**Figure 1.**
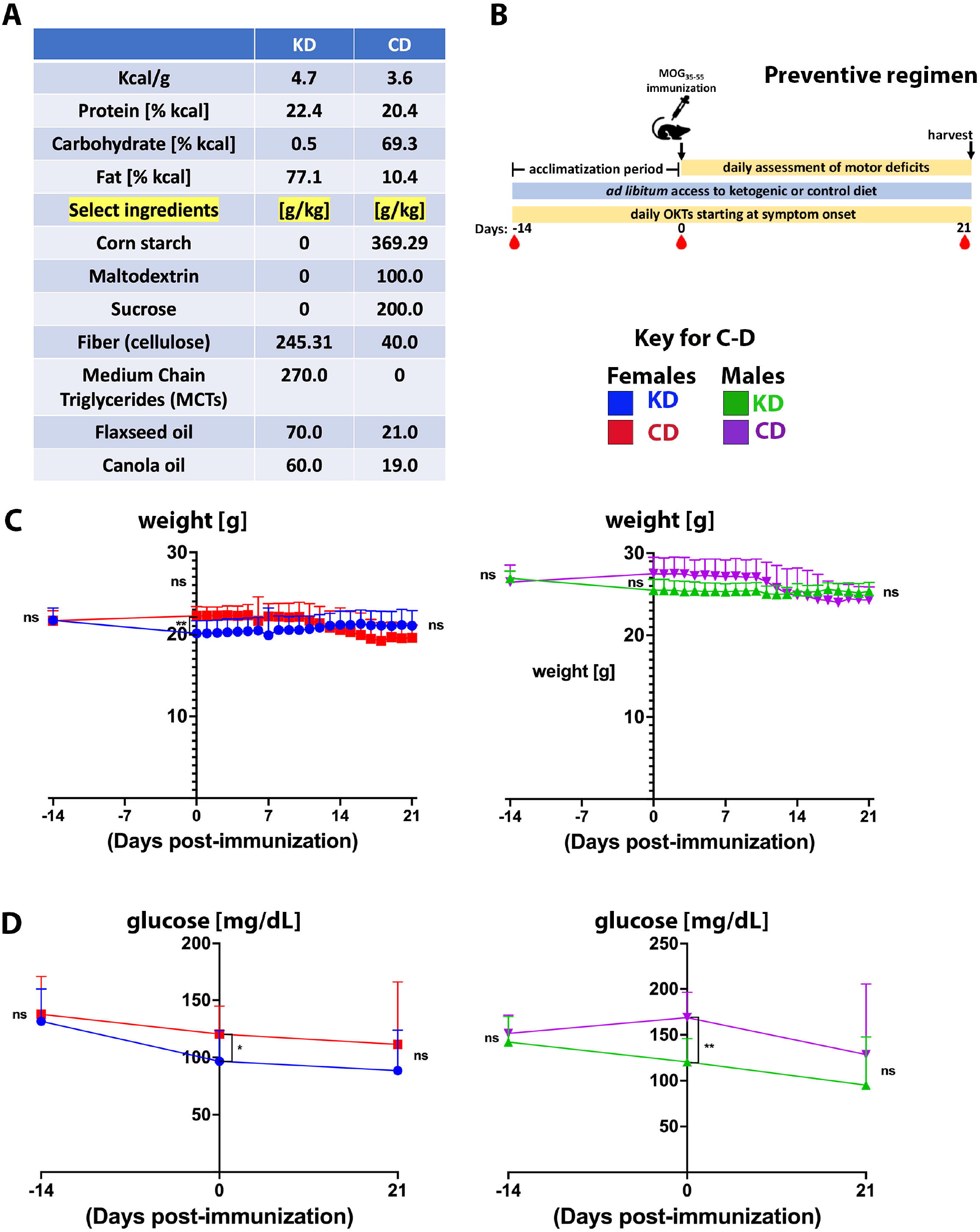
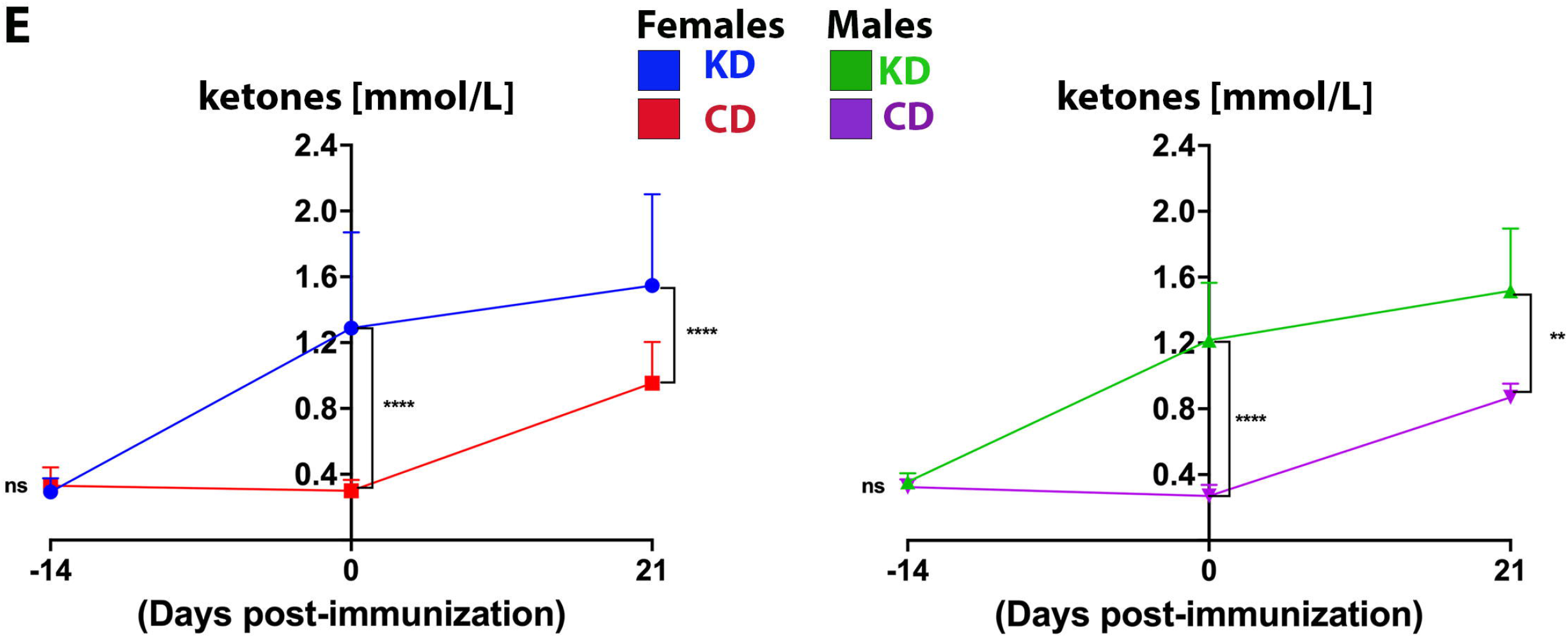
The KD stabilizes body weight and blood glucose but increases circulating ketones. **(A)** Macromolecular compositions of diets. The sources of carbohydrate in the CD are corn starch, maltodextrin, sucrose, and fiber in the form of cellulose. The only carbohydrate in the KD is cellulose. The fat content of the KD is derived from MCTs (C8 and C10), flaxseed oil, and canola oil. **(B)** Outline of experimental approach for the preventive regimen. Mice were fed a KD or CD for 2 weeks prior to MOG_35-55_ immunization to acclimate and were maintained on their respective diets for the duration of the experiment. Motor scores and visual acuity were tracked daily, and blood draws (red drop symbol) for glucose and ketone levels were taken at days −14, 0, and 21 dpi. Tissues and plasma were harvested 3 weeks post immunization at study termination. **(C-E)** Graphs of body weights (**C**), blood glucose levels (**D**), and blood ketones (**E**) as a function of days before and after MOG_35-55_ immunization. For graphs, blue and green traces show female and male mice on KD, respectively. Red and purple traces show female and male mice on CD, respectively. N=10-17 mice/sex. Whisker-bar standard deviations and p-values for differences between curves in C were computed with mixed linear model implementation of lme function, nlme R package. Asterisks denote statistical significance: * p<0.05, ** p<0.01, *** p<0.001, **** p<0.0001; ns = not statistically significant. Data compiled from 4-6 independent experiments.

### MOG-EAE

EAE was induced in 10-12 week old mice by subcutaneous flank injection of 150 μg of myelin oligodendrocyte glycoprotein peptide (residues 35-55; MOG_35-55_) (Genemed Synthesis, Inc., San Francisco, USA) emulsified in incomplete Freund’s adjuvant (Thermo Fisher Scientific; DF0639606) supplemented with 5 mg/mL heat-inactivated *Mycobacterium tuberculosis* (Thermo Fisher Scientific; DF3114338). Mice were injected intraperitoneally with 250 ng *Bordetella pertussis* toxin (List Biological Laboratories, Inc. #181) the day of and 2 days following MOG_35-55_ immunization. For the prevention experiments, mice were euthanized for post-mortem histology 21-22 days post-immunization (dpi). For the intervention experiments (Figure 7), all mice were maintained on standard chow until symptom onset at which time animals were switched to either the KD or CD and followed until 28 dpi. For the study presented in Supplemental Figures 7B-C, KD-fed mice were followed until 35 dpi. Manifestations of progressive ascending paralysis of classical EAE were assessed daily, using a more granular scoring system than we previously described (35–37): 0 – no disease, 0.5 – reduced tail tone, 1-loss of tail tone, 1.5 – limp tail and ataxia, 2-hind limb paresis, 2.5 – one hind limb paralyzed, 3-complete hind limb paralysis, 3.5 – complete hind limb paralysis and forelimb weakness, 4-hind limb paralysis and forelimb paresis, 5-moribund or dead. Mice were weighed daily to ensure weight loss did not exceed 25% of starting weight at the time of immunization.

**Figure 2.**
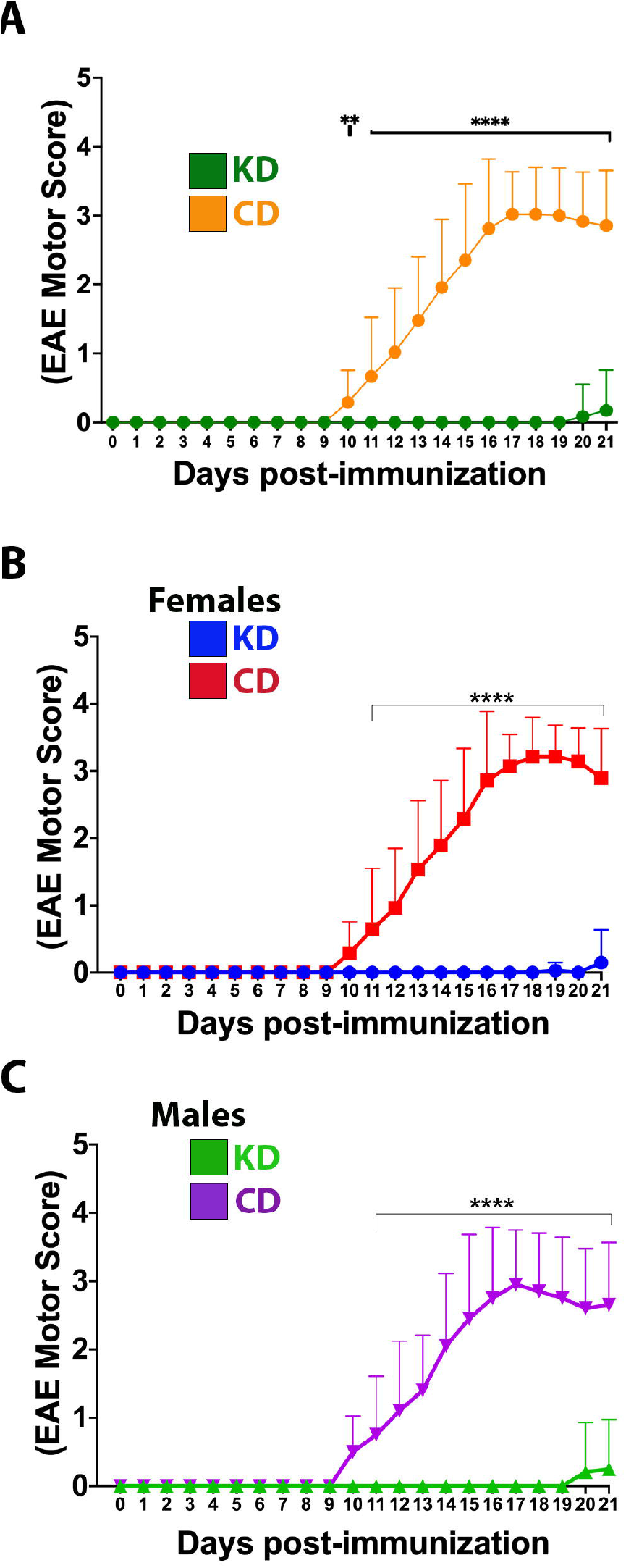
The KD prevents EAE-induced motor deficits. **(A)** Daily motor scores of EAE mice on the KD (green) versus the CD (orange). **(B)** Daily motor scores as a function of diet for female mice. Blue trace shows data for KD-fed mice and red trace shows data for CD-fed mice. **(C)** Daily motor scores as a function of diet for male mice. Green trace shows data for KD-fed mice and purple trace shows data for CD-fed mice. N=10-17 mice/sex/diet. Whisker-bar standard deviations and p-values for differences between curves were computed with mixed linear model implementation of lme function, nlme R package. Asterisks denote statistical significance: ** p<0.01, **** p<0.0001. All data compiled ≥ 4 independent experiments.

**Figure 3.**
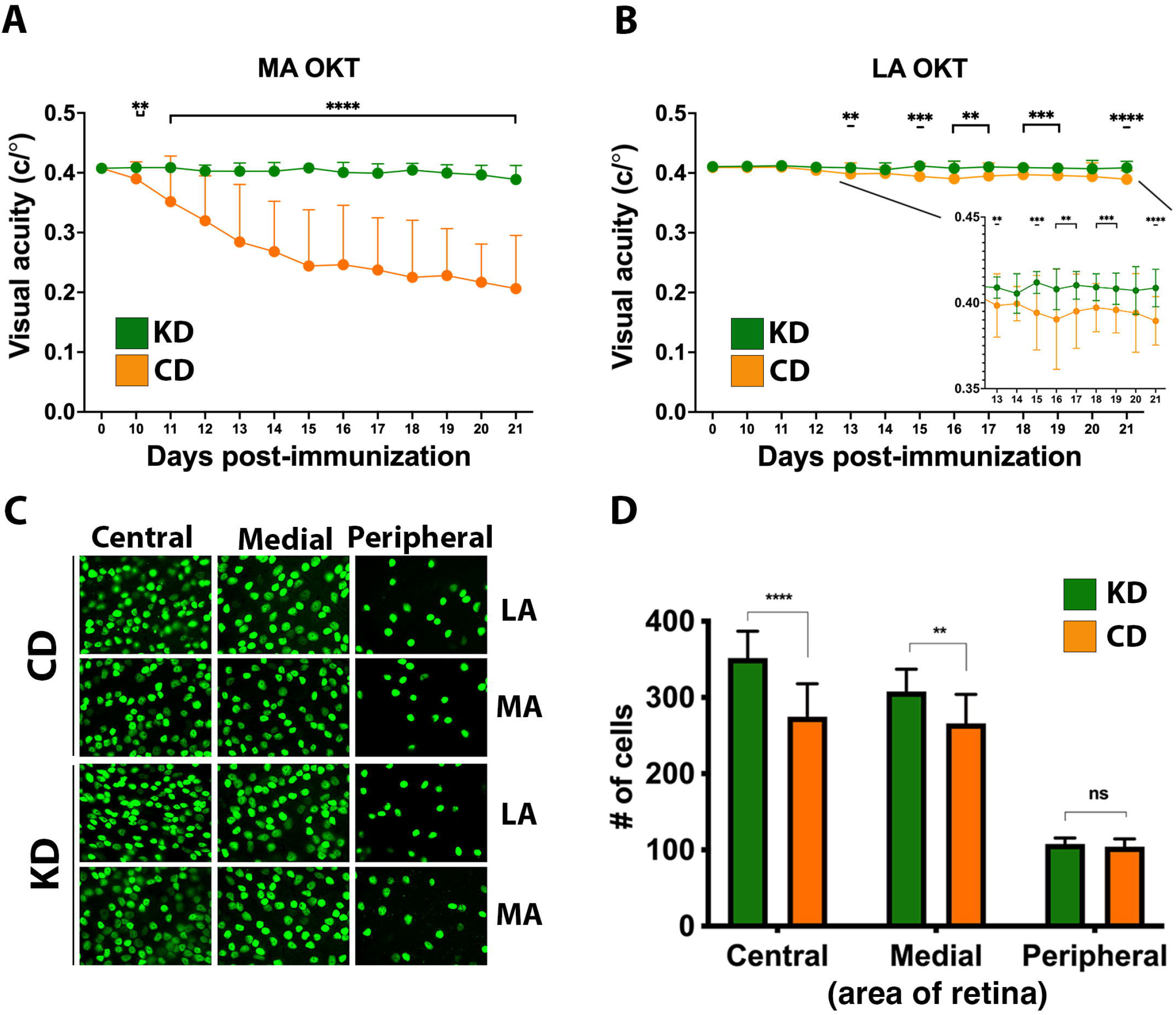
The KD prevents visual acuity deficits and spares retinal ganglion cells in the central retinas of EAE mice. **(A)** OKT measurements of visual acuity of the more affected (MA) eyes of EAE mice fed the KD (green) or CD (orange); n=14-17 mice/diet. **(B)** Same as (A) for less affected (LA) eyes; n=10-12 mice/diet. Inset with adjusted scales of x-axis and y-axis highlights significant differences in visual acuity between diets. Whisker-bar standard deviations and p-values for differences between curves in (A) and (B) were computed with mixed linear model implementation of lme function, nlme R package. **(C)** Representative photomicrographs of Brn3A^**+**^ RGCs in the central, medial, and peripheral retinas of mice as a function of diet. Examples are shown for both the MA and the LA eyes. **(D)** Graph of Brn3A^**+**^ RGC cell counts in central, medial, and peripheral retinas of MA eyes as a function of diet. Data are color coded based on diet as in (A) and (B). Data compiled ≥ 4 independent experiments.

**Figure 4.**
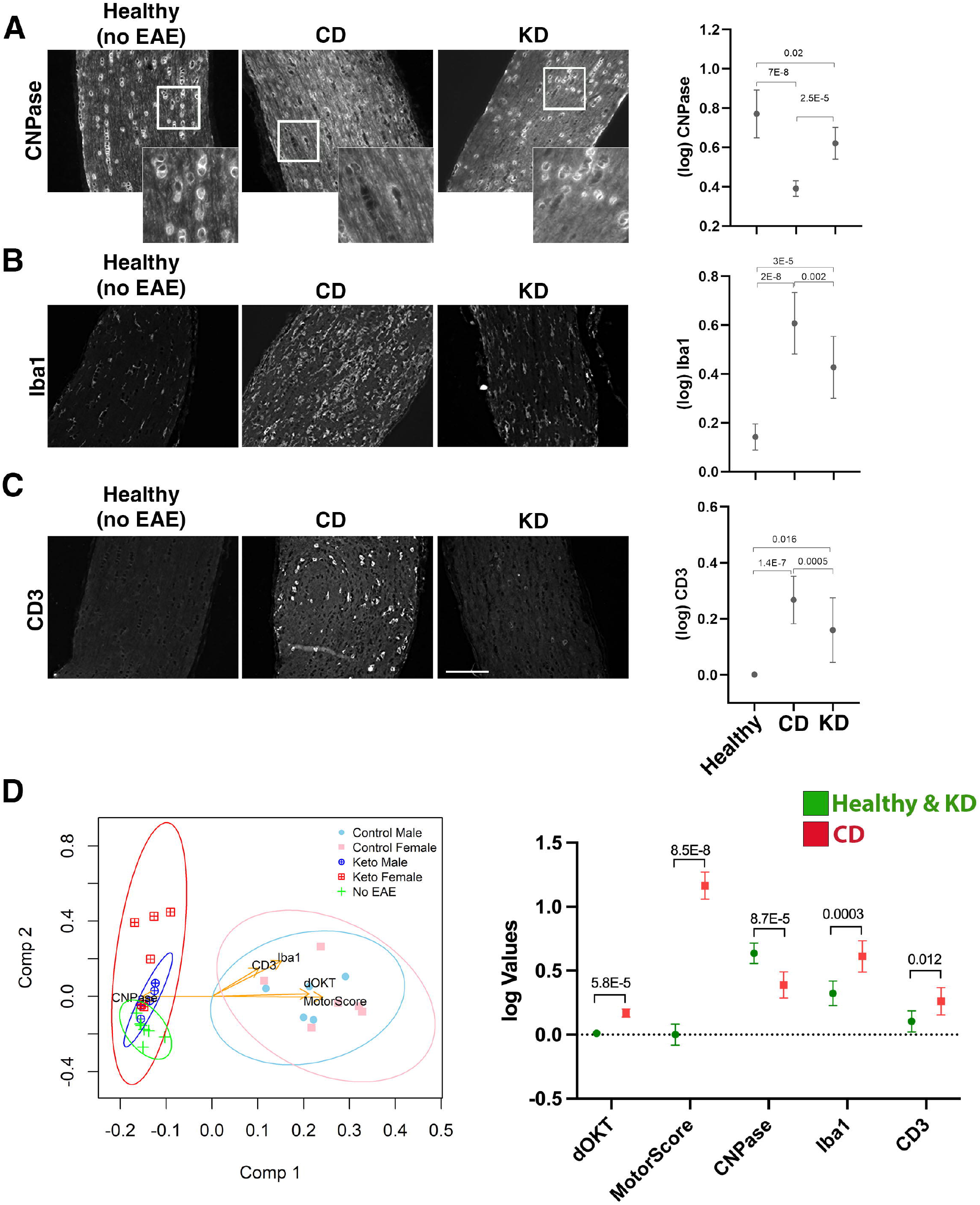
The KD preserves oligodendrocytes on the optic nerve and reduces immune cell infiltrates. **(A-C)** Representative photomicrographs of paraffin-embedded optic nerve sections labeled for the indicated markers (CNPase, Iba1, and CD3, A-C respectively). Optic nerves from the MA eyes of EAE mice consuming the indicated diets and from healthy (no-EAE) mice on standard chow were compared. EAE MA optic nerves were taken from the mice analyzed functionally by OKT in Fig. 3. Sections spanning the entire length of the optic nerve were labeled with each marker. Insets in (**A**) highlight the perinuclear CNPase labeling that was quantified. Mean values for CNPase, Iba1, and CD3 labeling of optic nerve sections are shown in the graphs to the right of the photomicrographs. Whisker bars depict 95% confidence intervals for each variable and horizontal bars with accompanying p-values above the bars indicate statistical significance as calculated by unequal group variance ANOVA with Tukey’s post hoc adjustment. Size bar in (**C**) corresponds to 100 *μ*m. **(D)** Principal components (PC) biplot to summarize, in 2d projection, the similarities among mice groups and their relationships to the 5 measured scores (i.e., motor scores, changes in OKT from baseline, CNPase, Iba1, CD3). The dots on the graph are samples, colored according to their phenotype group, with the measured variables shown as arrows. Arrow coordinates on the two axes show each variable contributing loading on the first two principal components. Multidimensional 95% confidence regions for each mouse group, projected in 2d, are shown as ellipses. All data acquired from n≥ 4 mice/sex/diet. The graph to the right of the PC biplot makes explicit, for the five markers, the difference between the two clusters of mice (Healthy+KD versus CD) separated by the first principal component. Whisker bars depicting 95% confidence intervals are shown for each variable both in Healthy+KD mice and in CD mice with the p-values assessing the significance of their differences marked on the graph.

**Figure 5.**
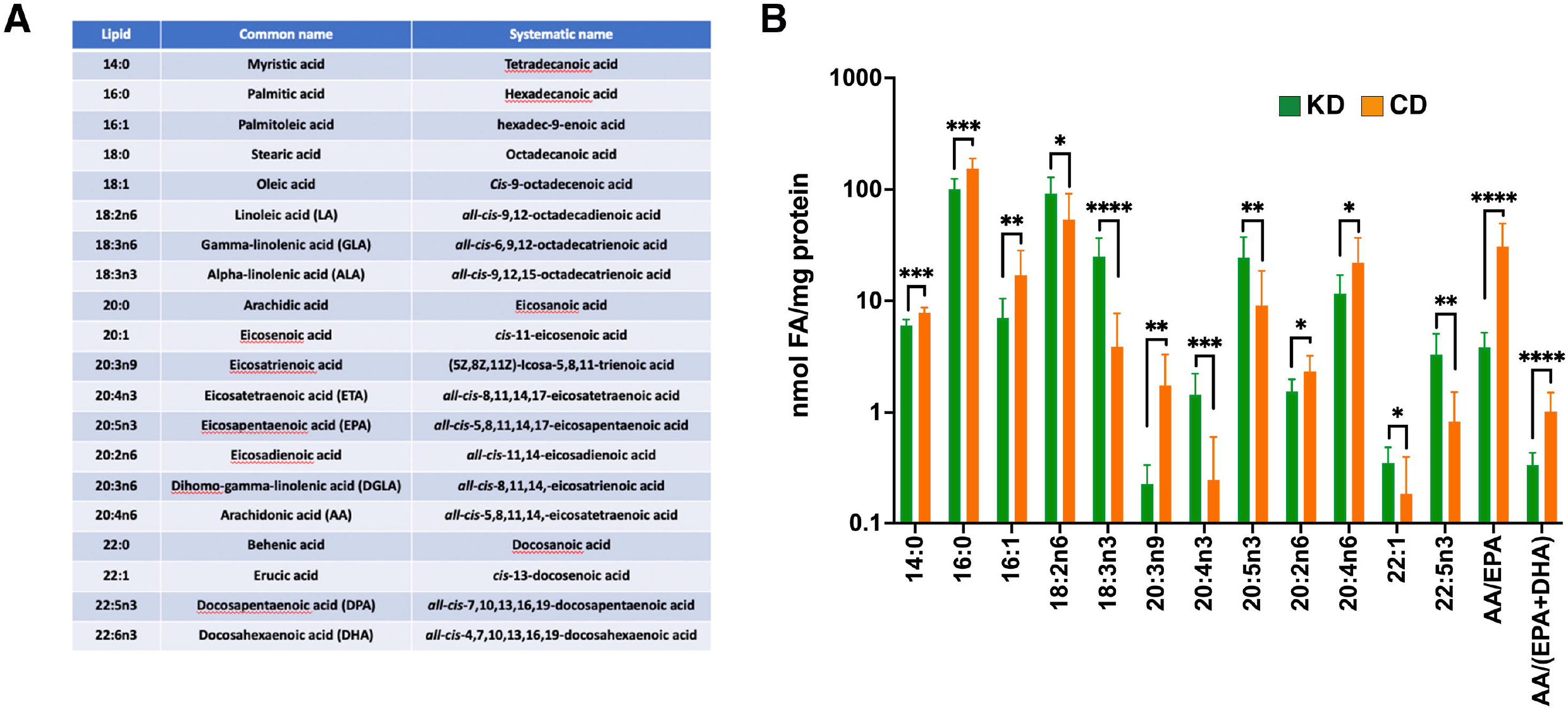
The KD increases plasma ω3 fatty acid content and suppresses levels of pro-inflammatory ω6 fatty acids. **(A)** List of fatty acids analyzed with the accompanying omega designations (aka. lipid), common names, and systematic names. **(B)** Graph depicting nmols of individual fatty acids per mg of protein in the plasma that are statistically significantly different (*p≤ 0.05, **p<0.01, ***p<0.001, ****p<0.0001 as determined by Mann-Whitney test with Bonferroni correction) between EAE mice fed the KD (green) versus the CD (orange). The ratio of arachidonic acid (AA; 20:4n6) to (EPA +DHA) (20:5n3 + 22:6n3) is included as a proxy of systemic inflammation. Note: graph uses log scale on the y-axis with data derived from n≥6 mice per diet from ≥2 independent experiments.

**Figure 6.**
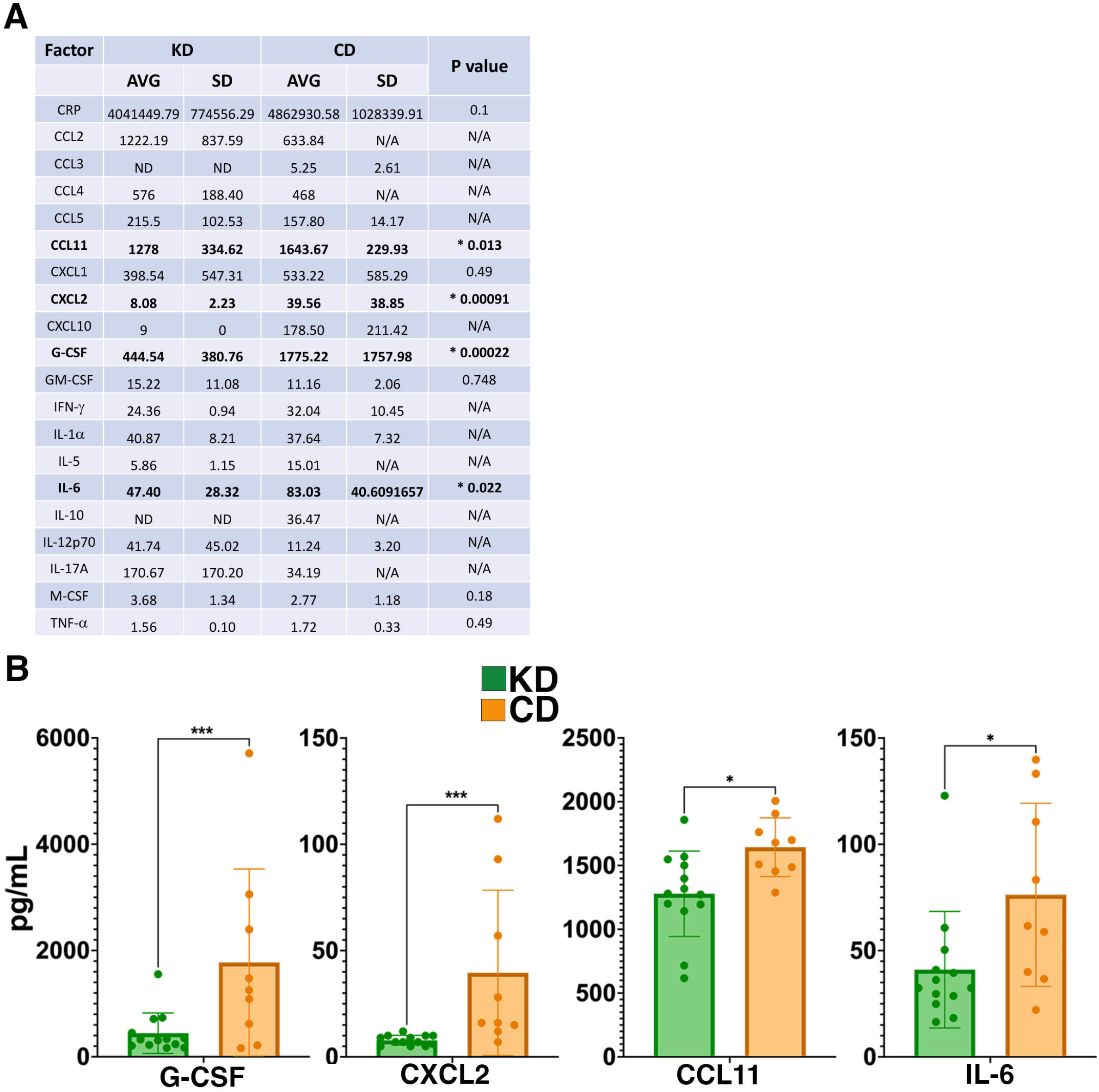
The KD suppresses levels of G-CSF, CXCL2, and other markers of inflammation. **(A)** Table of cytokine and chemokine measurements from the plasma of EAE mice. Asterisks and bolding highlight factors that are statistically significantly different as a function of diet. ‘AVG’ denotes average and ‘SD’ denotes standard deviation. ‘ND’ indicates that all samples analzyed were below the detection limit of the assay and ‘N/A’ indicates that insufficient numbers of samples were within the detectable range of the assay to perform statistical analyses. **(B)** Graphical presentation of the 4 factors that differ in the plasma 21 dpi as a function of diet. EAE mice consumed either the KD (green) or a CD (orange), 4-8 mice per diet per sex from ≥2 independent experiments. Statistical significance for cytokines and chemokines was determined with robust linear regression using the rlm function in MASS R package. p values relied on robust F-Testing (Wald) performed with f.robftest function from sfsmisc package. For graphs in (B), * p<0.05, *** p<0.001.

**Figure 7.**
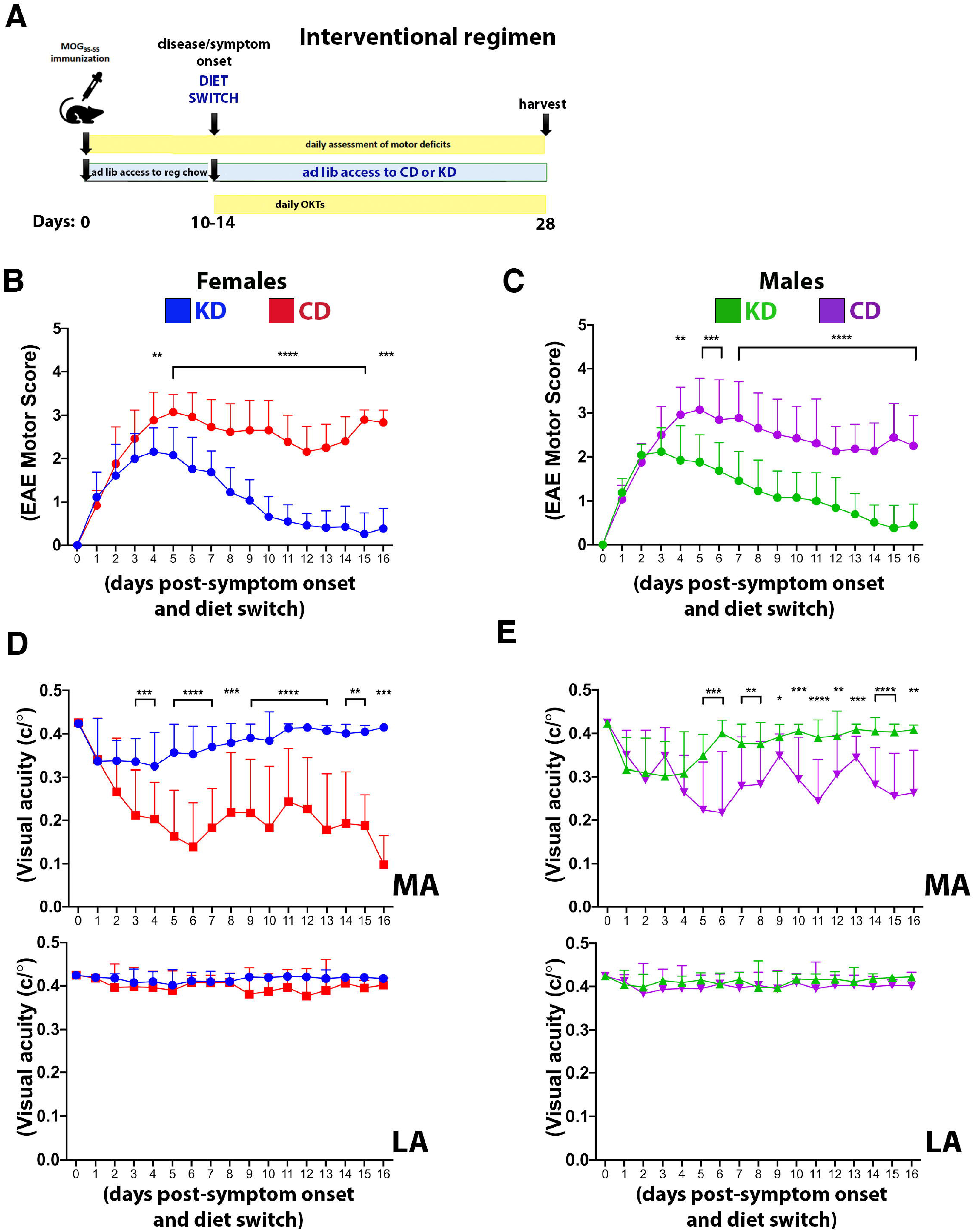
The KD restores motor and visual function to EAE mice when fed following symptom onset. (**A**) Diagram of the interventional regimen. Mice were immunized with MOG_35-55_ (day 0), switched to either KD or CD diets following symptom onset (10-14 dpi), and maintained on the respective diets for the duration of the experiment (28 dpi). Motor scores and visual acuity were tracked as indicated. Daily motor scores of female (**B**) and male mice (**C**) switched to the indicated diets following the onset of motor and visual deficits. Blue and green traces show female and male mice on KD, respectively, with red and purple traces showing female and male mice on CD, respectively. n=13 mice/sex/diet. **(D)** OKT measurements of visual acuity of the more affected (MA; top graph) and less affected (LA; bottom graph) eyes in female mice on the indicated diets. n=13 mice/diet. **(E)** Same as (D) for male mice. n=13 mice/diet. All data graphed with the x-axis representing “days post-symptom onset and diet switch”. N.B.: DAY 0 is the day *before* symptom onset and is denoted by a motor score of zero. Asterisks in all graphs denote statistical significance: * p<0.05, ** p<0.01, *** p<0.001, **** p<0.0001. Data compiled ≥ 4 independent experiments.

### Visual acuity assessment

Visual acuity threshold was measured daily by OKT response using Optometry software and apparatus (Cerebral Mechanics Inc., Alberta, Canada) as previously described (35–37). Briefly, mice are placed on a pedestal inside a box with a virtual cylinder consisting of vertical lines projected on 4 computer screens of the box surrounding the animal. The vertical lines rotate at varying frequencies, and tracking behavior is assessed in a stepwise manner as the thickness of the lines are reduced. Visual acuity is represented as the highest spatial frequency at which mice track the rotating cylinder. Optokinetic tracking is a temporal-to-nasal reflex, and therefore counter-clockwise and clockwise rotations exclusively test the right and left eye, respectively. When EAE-induced motor deficits rendered mice without the balance and stability to adequately perform OKT testing, a measurement was not recorded for the affected mouse on that day.

### Retinal flatmount analysis

Retinal flatmounts were prepared and RGCs labeled and quantified as described (35–37) using anti-Brn3a (goat; Santa Cruz, Santa Cruz, CA; sc31984; 1:500), Alexa546nm Fluor-conjugated donkey anti-goat IgG (Molecular Probes; 1:1,000), and Hoechst 33342 (2 μg/mL), all diluted in 3% BSA/10% donkey serum/PBS. Retinas were washed in PBS before mounting with Prolong Gold mounting medium (Life Technologies, Grand Island, NY) and examined with a Nikon 80i microscope with a 60X objective. Images were captured with a DXM1200C camera using NIS-Elements software (Nikon, Inc. Tokyo, Japan). Photomicrographs were captured from the four leaflets comprising the flatmount with representative images captured from the peripheral, medial, and central retina within each quadrant, yielding 12 pictures per retina. Images were identically contrast-enhanced. Brn3a-positive RGCs were counted manually using the FIJI Cell Counter Plugin.

### Fatty acid analysis

Plasma fatty acid profiles were determined by extracting total lipids from 40-50 μl of plasma following the method of Bligh and Dyer (38) with slight modifications (39). The purified total lipid extracts were stored under nitrogen until use. To these lipid extracts, 50 nmol each of 15:0 and 17:0 internal standards were added and the total extracts subjected to acid hydrolysis/methanolysis to generate fatty acid methyl esters (FAMEs) (40). All reagents for fatty acid extraction and derivatization were of the highest quality available from Sigma-Aldrich. FAMEs were identified using an Agilent Technologies 7890A gas chromatograph with a 5975C inert XL mass spectrometer detector (Agilent Technologies, Lexington, MA) as described (40). The gas chromatograph-mass spectrometer was operated in the electron impact total ion monitoring mode. The injection volume was 1 μl and the inlet, held at 325°C, was set to pulsed splitless mode. An Agilent Technologies HP-5MS column (30 m × 0.25 mm × 0.25 μm) was used with a helium carrier gas flow rate of 1.2 ml/min. The oven temperature began at 130°C for 1.0 min, was ramped to 170°C at 6.8°C/min, and was then ramped to 215°C at 2.9°C/min. After holding at 215°C for 15.0 min, the oven was ramped to 260°C at 20°C/min and held for 5.0 min. The oven was then ramped to 325°C at 15°C/min and held for 18.0 min. The mass spectrometer transfer line, ion source, and quadrupole temperatures were 325°C, 230°C, and 150°C, respectively.

FAMEs were quantified using an Agilent Technologies 6890N gas chromatograph with flame ionization detector (GC-FID) (41). Sample concentrations were determined by comparison to internal standards 15:0 and 17:0. The injection volume was 1 μl and the inlet, held at 290°C, was set to pulsed split mode (10:1 ratio). An Agilent Technologies DB-23 column (60 m × 0.32 mm × 0.25 μm) was used with a hydrogen carrier gas constant pressure of 13.1 psi. The oven temperature began at 130°C for 0.8 min, was ramped to 170°C at 8.2°C/min, and was then ramped to 215°C at 3.5°C/min. After holding at 215°C for 9.5 min, the oven was ramped to 230°C at 50°C/min, and was then held for 8 min. The oven was then ramped to 290°C at 12.0°C/min and was held for 12 min. The detector was held at 290°C. Data for the diets is represented as μg of each fatty acid per mg of diet, and data from plasma is represented as nmol of fatty acid per mg of plasma protein.

### Blood glucose and ketone measurements

Blood ketone levels (non-fasting) were measured with a Precision Xtra Blood Ketone Monitoring System and fasting blood glucose was measured after 6 hrs of food withdrawal using a True Metrix Blood Glucose Meter. Blood was drawn from the tail vein.

### Cytokine analyses

Twenty-six cytokines and chemokines in the plasma of EAE mice 21 dpi were assayed using custom xMAP multiplex cytokine panels from Biotechne, Inc. according to the manufacturer’s recommendations and processed in the OMRF Arthritis and Clinical Immunology Human Phenotyping Core. 6 factors were below the detection limit of the assay in all samples (IL-1β, IL-2, IL-4, IL-13, IL-17A, and IL-27) and excluded from the table in Figure 6.

### Immunohistochemistry of optic nerves

To assess oligodendrocytes (as a marker of myelination) as well as lymphocyte and macrophage infiltration, we incubated sequential de-paraffinized sections with anti-2’,3’-cyclic-nucleotide 3’-phosphodiesterase (CNPase) (mouse CNPase: 1:200 dilution; Biolegend, Inc.), anti-CD3 (rabbit; 1:250; Abcam) or anti-Iba1 (mouse; 1:200; Millipore, Inc.) antibodies after antigen retrieval in R-buffer B. Images were captured using a Nikon TE2000 fluorescent microscope. Quantification of staining was performed using FIJI Software. CNPase, Iba1, and CD3 staining on sections spanning the entire length of the optic nerve were quantified by a masked experimentalist using representative images from both ends and the middle of the optic nerve. The number of positive cells within a defined area were counted using the Cell Counter Plugin, and the number of positive-stained cells per 10,000 pixels was calculated for each optic nerve.

### Ex vivo stimulation of splenocytes

Twenty-one dpi, single cell suspensions from spleens were prepared. 2 x 10^6^ cells were plated in triplicate in complete RPMI (supplemented with 10% FBS, 2mM L-glutamine, 1% pen/strep, 2mM β-mercaptoethanol, non-essential amino acids, 1mM sodium pyruvate). 10μg/mL murine MOG_35-55_ peptide was added to the “stimulated” wells and incubated at 37°C for 24 hrs. For the final 5 hrs of the 24hr incubation, samples were spiked with 50ng/mL PMA (Sigma, Inc. cat # P1585), 500ng/mL ionomycin (Sigma, Inc. cat # IO634), and BD Golgi StopTM containing monensin (0.66μL/mL; BD Biosciences). Cells from triplicate wells were combined, stained with a fixable viability dye (Biolegend) and treated with Fc block (TruStain FcX PLUS; BioLegend, Inc. cat # 156604) subsequent to staining with the following anti-mouse Abs against cell surface markers (all Abs from eBioscience, Inc.): anti-CD3 (145-2C11), anti-CD4 (GK1.5), anti-CD8 (53-6.7), and anti-CD44 (IM7). Cells were fixed and permeabilized with BD Cytofix/CytopermTM Plus (BD Biosciences, Inc.) and stained with anti-IL17 (TC11-18H10.1) and anti-IFN-γ (XMG1.2) to detect the respective cytokines intracellularly. Samples were processed on a LSRII flow cytometer (BD Biosciences) and data analyzed with FlowJo version 10.7.1 software (BD Biosciences, Inc.).

### Statistical analysis

Confidence intervals and p values for the statistical significance of each studied effect in the longitudinal or binocular data were determined by fitting the data to a linear mixed-effects model, using the lme function implemented in the nlme R package, as detailed in (35, 42–44). This function is an extended version of regular linear regression but can accommodate complex data collection design features, such as longitudinal measurements, nested layers, and within-group correlation. Standard testing methods, such as the Student’s paired or unpaired t-test and the Mann–Whitney exact test, with Bonferroni correction for multiple testing, were employed when no embedding was involved. Graphpad Prism 9 was used to display the results. Testing for differences in marker levels among the five mice groups in Figure 4 was performed following log transformations for improving normality by unequal group variance ANOVA and Tukey’s post hoc adjustment. Differences in cytokine and chemokine levels in Figure 6 were assessed with robust linear regression using the rlm function in MASS R package, and p value computations relied on robust F-Testing (Wald) performed with f.robftest function from sfsmisc package. Principal component exploratory analysis (PCA) and biplot generation were carried out with specific functions from stats package in R. Multidimensional 95% confidence regions were added to the biplot using the draw.ellipse function from the plotrix package. Throughout the manuscript, the asterisks code denoting significance is: * p<0.05, ** p<0.01, *** p<0.001, **** p<0.0001; ns = not statistically significant.

## RESULTS

The current study tested the efficacy of a KD in the EAE mouse model of autoimmune-mediated demyelination. Many pre-clinical studies that have demonstrated therapeutic efficacy with a KD have used formulations containing lard, soybean oil, and hydrogenated fats (e.g., (22, 45, 46)). In contrast, the KD used here is enriched in fiber and the sources of fat are MCTs, flaxseed oil, and canola oil (Fig. 1A). This diet was designed to be anti-inflammatory by minimizing ω6 fatty acids and hydrogenated fats (19). A complete list of the diet ingredients is provided in Supplemental Fig. 1A.

The experimental paradigm used to test the KD in a preventive regimen is shown in Fig. 1B. Animals were fed either a KD or an ingredient-matched control diet (CD) for 2 weeks prior to MOG_35-55_ peptide immunization to acclimate mice to the diet based on previous strategies testing dietary impacts on CNS autoimmunity in the EAE model (e.g., (47). Mice were maintained on their respective diets for the duration of the experiment during which motor scores, vision, and body weights were longitudinally tracked and blood was collected. 3 weeks post-immunization, mice were euthanized for post-mortem analyses. Notably, consumption of the KD for 5 weeks in the absence of EAE led to modest weight reduction whereas 5 weeks of the CD did not (Supplemental Fig 1B). Animals were randomized to KD or CD groups and had comparable weights within each sex at study initiation (Fig. 1C). From immunization to completion of experiments at 21dpi, among EAE mice fed the KD, females slightly gained weight whereas males were weight stabilized. In contrast, both sexes on the CD undergoing EAE lost body weight (Supplemental Fig. 1C), consistent with sickness behavior (e.g., inflammatory anorexia) (48). Compared to the CD, the KD reduced fasting blood glucose levels after 2 weeks of feeding despite no differences at study onset or termination (Fig. 1D). Elevated non-fasting ketones are a defining molecular signature of a KD (49) and circulating, non-fasting ketones (β-hydroxybutyrate specifically) were increased in mice fed the KD for 2 weeks and persisted through the duration of the study (Fig. 1E). Mice on the CD also had moderately elevated circulating ketones three weeks after immunization, likely related to sickness behavior (e.g., inflammatory anorexia) (48). These data show that the KD lowered fasting blood glucose and induced nutritional ketosis but did not promote obesity during 5 week-long experiments.

### KD PREVENTS THE ONSET OF MOTOR AND VISUAL DEFICITS IN EAE MICE

Remarkably, mice fed the KD were spared the ascending paralysis and motor deficits induced by EAE (Fig. 2A), with comparable efficacy observed for females and males (Figs. 2B and 2C, respectively). Because optic neuritis and visual disturbances are major sequelae of autoimmune demyelinating disease in humans (50–52), we complemented the motor score measurements with visual acuity measurements using optokinetic tracking (OKT). OKTs were recorded at the initiation of the diets, prior to MOG_35-55_ immunization, and then daily beginning ~10 dpi, coincident with symptom onset in CD-fed animals (Fig. 1B). As we reported previously (36), mice subjected to MOG-EAE typically undergo episodic *monocular* vision loss, consistent with what is experienced by people with relapsing-remitting MS (RRMS) (53, 54). These deficits are readily detectable as diminished visual acuity in one eye (35, 36). The OKT measures each eye separately, so every mouse has a designated more affected (MA) and less affected (LA) eye. Coincident with the mitigation of motor deficits, the KD preserved vision in the MA eyes of EAE mice (Fig. 3A). Although visual deficits were minimal in the LA eyes of mice on either diet, the LA eyes of KD-fed mice retained better vision than their CD-fed counterparts (Fig. 3B, inset). In alignment with the efficacy of the KD on motor function, this preservation of vision was independent of sex (Supplemental Figs. 2A-B).

The axons of retinal ganglion cells (RGCs) are bundled together into the optic nerve and episodes of optic neuritis induce RGC apoptosis (55–59). We therefore quantified RGC counts and found that the functional preservation of vision mediated by the KD was accompanied by a sparing of Brn3A^+^ RGCs in the central and medial retina of MA eyes from EAE mice (Figs. 3C-D). As we reported previously (36), RGCs in the peripheral retina are not typically lost during MOG-EAE (Fig. 3D), fitting with RGC loss in longstanding MS cases being most prominent in the central retina, nearest the optic nerve head (60). When analyzed as a function of diet and sex, the KD-mediated sparing of RGCs in the MA eyes was significant in the central retina for both females and males (Supplemental Fig. 2C).

### THE KD PRESERVES OLIGODENDROCYTES AND RESTRICTS INFLAMMATORY INFILTRATION OF THE OPTIC NERVE IN EAE MICE

To complement the OKT and RGC analyses, immunohistochemistry (IHC) was performed on paraffin-embedded optic nerves from the MA eyes of the mice analyzed for visual function in Fig 3. For each marker, labeling was quantified from sections captured along the entire length of the optic nerve, and optic nerves from healthy (i.e., no EAE) mice were included for comparison. CNPase, an oligodendrocyte marker and proxy of myelination, was largely preserved on the optic nerves of KD-fed mice (Fig. 4A). CNPase labeling was quantified by comparing perinuclear staining, with the specificity of this staining corroborated by labeling optic nerves from healthy mice (Fig. 4A, left panel inset). Labeling for Iba1 and CD3 to mark macrophages/microglia and T cells, respectively, revealed that mice fed the KD had reduced Iba1 and CD3 labeling of the optic nerve compared to mice fed the CD (Figs. 4B and 4C, respectively).

When the data were further examined to determine the combined impacts of diet and sex, both females and males on the KD had significantly preserved CNPase labeling compared to their CD-fed counterparts (Supplemental Fig. 3A). Likewise, KD-fed male optic nerves had decreased Iba1 and CD3 labeling compared to CD-fed males (Supplemental Figs. 3B and 3C, respectively). Due to higher variability between samples, differences in Iba1 and CD3 labeling did not reach significance between the KD and CD female mice (Supplemental Figs. 3B and 3C, respectively), despite the preservation of visual acuity for female mice on the KD (Supplemental Fig. 2A). Notably, all groups had statistically significant increases in Iba1 labeling compared to optic nerves from healthy mice (Supplemental Fig. 3B), despite no loss of visual acuity for the KD group (Fig. 3A). For CD3 labeling, females and males on the KD were not statistically different than healthy mice whereas EAE males and females on the CD showed significant increases in CD3^+^ infiltrates (Supplemental Fig. 3C).

Principal components exploratory analysis (Fig. 4D) confirmed that the relatively higher CNPase labeling and lower Iba1 and CD3 labeling for mice fed the KD correlated with lower motor and visual acuity deficits. The first principal component reveals that the KD-fed mice are statistically comparable to healthy mice but significantly different from the CD-fed group, presenting significantly improved motor and visual acuity scores and higher CNPase levels, along with lower Iba1 and CD3 levels. The second component suggests marginal separation of the healthy mice group from KD-fed mice, in terms of Iba1 and CD3 levels.

### A KD INCREASES CIRCULATING ω3 FATTY ACIDS IN EAE MICE

To test the hypothesis that the therapeutic efficacy of the KD derives from inducing a systemic anti-inflammatory milieu, we analyzed a panel of circulating fatty acids (FAs) in the plasma of EAE mice 21 dpi (Fig. 5A). With a focus on those FAs showing statistically significant differences between diets, we observed that, compared to the CD, the KD elevated circulating levels of 18:3n3, 20:4n3, 20:5n3, and 22:5n3 whereas the saturated fats, 14:0 and 16:0, the mono-saturated fat 16:1, and 20:4n6 (arachidonic acid; AA) were all decreased (Fig. 5B). Likewise, the ratio of 20:4n6 to (20:5n3+22:6n3) (i.e., AA/(EPA+DHA)), a proxy of inflammatory status, was decreased by the KD (Fig. 5B). Notably, 20:5n3, 22:5n3, and 22:6n3 are biosynthetic precursors of E-series, T-series, and D-series resolvins, respectively, specialized pro-resolving lipid mediators (SPMs) that restrict the development of chronic inflammation by dampening acute inflammation (reviewed in (61)). Consistent with the KD limiting systemic inflammation, 20:3n9 (eicosatrienoic acid; ETA) levels were decreased (Fig. 5B). This ω9 fatty acid can be converted to the proinflammatory C3 and D3 leukotrienes (62).

We also observed several circulating FAs changed as a combined function of sex and diet. Females fed the KD had increased serum levels of 18:2n6, 20:1, and 22:1 along with decreases in 16:1, 18:1, 20:2n6, and 20:4n6 (Supplemental Fig. 4A). EAE male mice fed the KD had elevated levels of the ω3 fatty acid 20:5n3 (Supplemental Fig. 4B), the intermediate in the conversion of 20:4n3 to 22:5n3 (63). 20:5n3 was also increased in KD-fed females but did not reach significance because of high variability among 20:5n3 levels within female CD-fed mice (data not shown). Together, these data are consistent with the KD conferring protection from EAE pathologies at least in part by enriching the systemic milieu with ω3 fatty acids associated with SPM-mediated neuroprotection and decreasing fatty acids, such as 20:3n9, that are precursors for the biosynthesis of pro-inflammatory leukotrienes.

To determine which of the enriched circulating ω3 fatty acids in EAE mice consuming the KD were contributed directly by the diet, we analyzed the fatty acid content of each diet (Supplemental Fig. 4C). Notably, compared to the CD, the additional flaxseed oil in the KD led to an enrichment by 4.3 *μ*g fatty acid/mg sample (or 1.6-fold) in 18:3n3 (a-linolenic acid; ALA). 18:3n3 can be biosynthetically converted *in vivo* to the corresponding 20:4n3, 20:5n3, and 22:5n3 (63, 64). As 20:4n3, 20:5n3, and 22:5n3 were absent from the KD and CD diets, the elevated levels of these ω3 fatty acids in the plasma of EAE mice consuming the KD (Fig. 5B) appear to be derived from the endogenous conversion of 18:3n3.

### CYTOKINES IMPACTED BY THE KD IN EAE MICE

Complementary evidence supporting the anti-inflammatory milieu resulting from the KD came from an analysis of 26 different cytokines and chemokines in the plasma 21dpi, of which 20 were detectable in some or all samples within the sensitivity of the multiplex assay (Fig. 6A). Of the cytokines detected, granulocyte-colony stimulating factor (G-CSF), C-X-C Motif Chemokine Ligand 2 (CXCL2), C-C motif chemokine ligand 11 (CCL11), and IL-6 were all significantly decreased 21 dpi in the plasma of EAE mice consuming the KD (Fig. 6B). Two sex-specific changes were also detected (Supplementary Fig. 5). EAE females on the KD had elevated monocyte-colony stimulating factor (M-CSF) whereas males had reduced C-reactive protein (CRP). Curiously, CRP levels in KD female mice were comparable to the levels in KD-fed males but CRP was not elevated in the female CD-fed group (Supplementary Fig. 5).

### EAE MICE FED A KD MOUNT A T-CELL RESPONSE TO MOG_35-55_ IMMUNIZATION

To rule out that the protection conferred by the KD in the preventive regimen was not a consequence of the diet blunting an immune response to the MOG antigen, splenoctyes were isolated 21 dpi from EAE mice on each diet, stimulated *ex vivo* in culture with MOG_35-55_ peptide, and subsequently with PMA, iononmycin, and monensin prior to antibody labeling to identify activated T cells (i.e., CD3^+^, CD4^+^, CD44^++^) and their respective intracellular levels of IL-17 and IFN-γ by flow cytometry (Supplemental Fig. 6A). IL-17 and IFN-γ expression by MOG_35-55_-activated splenic T cells is a signature EAE response following immunization with MOG (e.g., (65, 66)). For comparison purposes, unstimulated sets of splenocytes from mice on either the KD or CD were similarly stained for intracellular IL-17 and IFN-γ. These analyses showed that the percent of live CD3^+^CD4^+^ CD44^++^ T cells expressing IL-17 in the spleen was comparable between diets whereas a greater faction of stimulated splenocytes from KD-fed animals expressed IFN-γ versus their CD-fed counterparts (Supplemental Figs. 6A and 6B). Notably, the level of activated CD3^+^CD4^+^ CD44^++^ T cells was similar between the KD and CD groups (Supplemental Fig. 6C).

Of note, we observed a small of percentage of “breakthrough” mice consuming the KD that began to exhibit EAE symptoms ~ 19-21 dpi (Supplemental Fig. 7A). To determine whether all or most KD-fed mice would manifest EAE symptoms if the experimental time course was extended, we tracked EAE mice on the KD for an additional 2 weeks (i.e., 2 weeks pre-immunization plus 5 weeks post-immunization). Compared to the 90% of CD-fed mice (100% males and 80% females) that showed symptoms within 10-16 dpi (11-12dpi for males and 10-16dpi for females), extending the study out to 7 weeks total revealed that disease incidence among KD-fed mice was ~ 33% (20% for females and 50% for males) with symptoms not detected until 22-33 dpi (22-33 dpi for males and 31-32 dpi for females) (Supplemental Fig 7B). Furthermore, the severity of motor deficits among the subset of KD-fed animals that eventually developed symptoms was significantly reduced compared to their CD-fed counterparts (Supplemental Fig. 7C). Together, these lines of evidence support that although most mice on the KD are protected from the overt pathologies induced by the EAE model, the underlying mechanism is not a failure to mount a T cell response to immunization with the MOG antigen. Collectively, the KD effectively prevented disease onset in most mice and mitigated disease severity in the subset of mice that manifested functional deficits.

### THE KD AS AN INTERVENTIONAL REGIMEN RESTORES MOTOR AND VISUAL FUNCTION

Complementary studies in female and male mice were performed to determine the efficacy of the KD as an interventional regimen (Fig. 7A). All mice were maintained on standard chow until EAE symptom onset. At the first observable sign of motor and visual deficits, mice were switched to either the KD or the CD for the remainder of the study. Within 4 days of consuming the KD, motor (Figs. 7B-C) and visual (Figs. 7D-E) deficits were significantly mitigated in both sexes and continued to improve such that motor and visual function were restored to near baseline levels by the termination of the study. Notably, these studies spanned 26-30 days from the day of MOG immunization until termination but the data are graphed with the x-axis representing “days post-symptom onset and diet switch” to normalize for all mice not manifesting their respective initial functional deficits on the same day. These results complement the above prevention studies and importantly, confer translational relevance and feasibility to this dietary strategy.

## DISCUSSION

Previous studies established that dietary interventions including caloric restriction (20), a fast-mimicking diet (21, 22), intermittent fasting (23, 24), and a KD (22, 25, 26) can ameliorate EAE motor and cognitive deficits in EAE mice. Our data expand on these findings by demonstrating that a KD preserves visual acuity (Fig. 3) and myelination of the optic nerve, reduces Iba1^**+**^ and CD3^**+**^ infiltrates (Fig. 4), and spares RGCs (Fig. 3). We further show that the benefits extend comparably to female and male mice and that the KD increased circulating levels of the anti-inflammatory ω3 fatty acids 18:3n3, 20:4n3, 20:5n3, and 22:5n3 while decreasing the pro-inflammatory fatty acids 20:3n9 and 20:4n6 (Fig. 5). We also detected a KD-mediated decrease in circulating cytokines implicated in EAE and MS, namely G-CSF, CXCL2, CCL11, and IL-6 (Fig. 6B). When tested as an intervention, the KD promoted rapid and nearly complete recovery of motor and visual function (Fig. 7). This finding is particularly promising for establishing the clinical feasibility of acutely implementing the KD in response to symptom relapse.

The diet composition of the KD used in the present study (19) differs with previous EAE studies that used KDs containing butter, corn oil, and lard (e.g., (22, 26)), sources of fat associated with hepatic dysfunction and inflammation (67, 68). The different KDs were efficacious in mitigating EAE sequelae, but a consideration of the cardiometabolic health risks of chronically consuming high fat diets is of relevance to people with MS as these individuals will likely implement long-term nutritional interventions. In this regard, our studies support that the use of a KD enriched in ω3 fatty acids including a-linolenic acid (ALA, 18:3n3) can elicit therapeutic benefits without promoting weight gain or increasing inflammation (Supplemental Fig. 1 and Fig. 6).

Because different KD formulations mitigate EAE pathologies despite containing various pro-and anti-inflammatory fats (22, 26), the therapeutic efficacy of KDs may derive from the production of ketones and/or the dramatic limitation of simple sugars and starch consumption common to these regimens. Consistent with these hypotheses, high dietary glucose increases disease severity in the EAE model by promoting T_H_17 cell differentiation via ROS-mediated TGF-β activation (69). In clinical work, a KD improved glycemic control and reduced the medications of people with obesity and diabetes more effectively than a 55% carbohydrate low glycemic index diet (70). For people with (pre)diabetes, a KD reduced glycosylated hemoglobin (HbA1C), fasting glucose, fasting insulin, weight, blood pressure, triglycerides, alanine aminotransferase, and high-density lipoprotein, consistent with safety and tolerability for long-term adherence. Remarkably, 53% of enrolled participants achieved disease resolution (30). Individuals that are overweight and implemented a KD also had decreased body weight, insulin resistance, and serum markers of inflammation (e.g., TNF-α, IL-6, IL-8, MCP-1) (71–73). These normalizing effects on insulin sensitivity and insulin resistance benefit people with metabolic syndrome and diabetes (reviewed in (74)) and likely extend to people with MS, as insulin resistance is associated with elevated disability scores (16, 17). Clinical trials with small cohorts of people with MS have also shown promising responses using (modified) paleolithic diets that eliminate or dramatically reduce the consumption of simple carbohydrates, ultra-processed foods, and other putative disease aggravators (e.g., gluten, dairy, legumes) (13, 75–81).

A potential mechanistic contribution mediating the efficacy of the KD in the present study comes from the anti-inflammatory actions of resolvins, a family of SPMs derived from ALA. In alignment with this idea, elevated dietary consumption of 18:3n3 has been inversely linked with the risk of developing MS (82). ALA was enriched in the KD (Supplemental Fig. 4C) as well as in the plasma of EAE mice fed the KD (Fig. 5B). Moreover, the ω3 fatty acids in the plasma of mice fed the KD may directly contribute to lower levels of inflammatory infiltrates in the MA optic nerves (Fig. 4) as ω3 fatty acids enhanced lesion recovery by decreasing phagocytic infiltration to the corpus callosum following demyelination (83).

The plasma of KD EAE females and males was enriched in ETA (20:4n3) and DPA (22:5n3) (Figure 5 and Supplemental Figs. 4A and 4B) and males were also enriched with EPA (20:5n3) (Supplemental Fig. 4B). 20:5n3 and 22:5n3 are intermediate biosynthetic precursors of E-series and T-series resolvins, respectively. DHA (22:6n3) and arachidonic acid (20:4n6), the intermediates from which the Resolvin D-series and the lipoxins are synthesized, respectively, were not increased and arachidonic acid levels were reduced. However, the lack of elevated DHA and arachidonic acid does not necessarily preclude contributions from their respective SPMs as efficient synthesis reactions could limit accumulation of these intermediates. A study using the MOG-EAE model and standard chow over a 45-day course showed PUFA metabolism is compromised during disease progression and that daily administration of exogenous Resolvin D1 decreased EAE pathologies (84). Clinical work has identified imbalances in pro-inflammatory eicosanoids and SPM levels in the plasma of people with MS as a function of disease progression and severity. The expression of multiple biosynthetic enzymes and receptors for SPMs were impaired in peripheral blood mononuclear cells from people with MS and their monocytes were less responsive to SPMs in culture (85). Such deficiencies in the SPM synthetic machinery may underlie the observation that elevated EPA in the serum of people with MS tracks with increased severity on the expanded disability status scale (EDSS). This study additionally reported that circulating arachidonic acid levels associate with relapse-free status (86). Notably, study participants were taking IFN-β or other disease-modifying therapies, which confounds comparing these data directly to our study or other stand-alone dietary intervention studies.

Fitting with the KD promoting a systemic anti-inflammatory milieu, we detected reductions in four pro-inflammatory cytokines and chemokines in the circulation as a function of diet: G-CSF, CXCL2, CCL11, and IL-6 (Fig. 6). G-CSF and CXCL2 were the two pro-inflammatory cytokines reduced in both sexes fed the KD (Supplemental Fig. 5). G-CSF levels are typically kept low but rapidly increase in response to stress and inflammation to stimulate the production and maturation of granulocytes and neutrophils. G-CSF governs early signaling necessary for EAE disease induction (87) with the neutrophils produced driving multiple steps of EAE and MS progression (e.g., (88–92)), fitting with the KD reducing G-CSF in the circulation and blocking symptom onset.

CXCL2 levels were also reduced by the KD (Fig. 6B). This factor is also known as macrophage inflammatory protein 2-α (MIP2-α), is produced by macrophages and neutrophils at sites of inflammation, and functions to recruit neutrophils during inflammation (93). Transient receptor potential melastatin 2 knockout mice also have reduced CXCL2 and a suppression of neutrophil infiltration into the CNS during EAE (94). Thus, reduced levels of circulating CXCL2 are consistent with disease mitigation (95–97) and fit logically with lower G-CSF levels limiting neutrophil maturation.

KD fed mice had reduced CCL11, also known as eosinophil chemotactic protein (eotaxin-1) (Fig. 6B). This chemokine is a putative biomarker of disease duration in people with secondary progressive MS (98). Curiously, in a rat EAE model, increased CCL11 expression was associated with a TH2 anti-inflammatory response but did not correlate with eosinophil recruitment. Elevated CCL11 was detected in the CSF and lymph nodes (but not in the serum) and correlated with decreased ED1^+^/Iba1^+^ macrophages in the spinal cord and with protecting the integrity of the blood-brain barrier (99). In contrast, CCL11 was found to be elevated in the spinal cords of mice undergoing MOG_35-55_ EAE (100). As CCL11 has been implicated in systemic inflammation and as a pathogenic factor in a range of neurodegenerative and neuroinflammatory diseases (reviewed in (101)), reduced CCL11 in the serum by the KD is consistent with an anti-inflammatory milieu.

The KD reduced circulating levels of IL-6 (Fig. 6B), a pro-inflammatory cytokine secreted by astrocytes, macrophages, and microglia as well as other cell types in the CNS (e.g., neurons and endothelial cells) (102). IL-6 is detectable in brain lesions (103) and in the CSF from people with MS (104) and plays critical roles in MS and EAE pathophysiology that include compromising blood-brain barrier integrity in combination with IL-17A (105), cooperating with transforming growth factor-β to drive the differentiation and expansion of auto-reactive T_H_17 cells (106–109), and damaging myelin (102). Reduced circulating IL-6 by a KD has been reported (e.g., (110–116)) although studies in select populations (e.g., (117, 118) and some mouse models have also reported that a KD may not change or may even increase levels of this cytokine (e.g., (119)). Reduced levels of IL-6 in the serum of KD-fed EAE mice are consistent with this dietary approach blunting CNS disease burden and maintaining motor and visual functions.

In conclusion, our work demonstrates the efficacy of a KD to preserve motor and visual function in mice undergoing autoimmune demyelinating disease. This dietary strategy limits systemic inflammation by reducing key cytokines involved in mediating the infiltration, activation, and differentiation of auto-reactive T cells and neutrophils into the CNS. We further posit that this KD provides an abundance of ω3 fatty acids for SPM biosynthesis, the products of which restrict acute inflammatory responses to self-antigens from transitioning to chronic inflammation and tissue damage. The observation that initiating the KD beginning at the time of symptom onset can resolve both motor and visual deficits (Fig. 7) supports the potential of this diet for direct translational application and improved patience compliance, a current barrier for nutrition-based therapeutic strategies (120–122).

## ACKNOWLEDGEMENTS

Research reported in this publication was in part supported by a Presbyterian Health Foundation Collaborative Sciences Grant to SMP and SK and in part by the National Eye Institute of the National Institutes of Health under Award Number R01EY033782. The content is solely the responsibility of the authors and does not necessarily represent the official views of the National Institutes of Health. We are grateful to members of the Plafker, Axtell, and Kovats laboratories at OMRF and the Agbaga lab at DMEI for helpful input and thank Jocelyn Washington-McCoy for masked quantification of histology slides. We thank Drs. Joel and Carla Guthridge for assistance with the multiplex cytokine studies. KZJ is grateful for the support of an OMRF pre-doctoral fellowship endowed by the Barrett Scholarship Fund. We also acknowledge unrestricted grant support from Research to Prevent Blindness and P30EY021725 to the Dean McGee Eye Institute for support of the Lipidomics core.

## AUTHOR CONTRIBUTIONS

**KZJ:** experimental design, acquisition and analysis of data, figure preparation, writing and editing. **DAW:** acquisition and analysis of data, editing. **KSP:** acquisition and analysis of data, figure preparation, writing and editing. **SK:** acquisition and analysis of data, editing. **CG:** statistical analyses, editing. **RSB:** acquisition and analysis of data, editing. **MT:** acquisition and analysis of data, editing. **MPA:** acquisition and analysis of data, editing. **SMP:** experimental design, figure preparation, writing and editing.

## REFERENCES

1. Leray E, Moreau T, Fromont A, Edan G. Epidemiology of multiple sclerosis. Rev Neurol (Paris). 2016;172(1):3–13.

2. Tullman MJ. Overview of the epidemiology, diagnosis, and disease progression associated with multiple sclerosis. Am J Manag Care. 2013;19(2 Suppl):S15–20.

3. Walton C, King R, Rechtman L, Kaye W, Leray E, Marrie RA, et al. Rising prevalence of multiple sclerosis worldwide: Insights from the Atlas of MS, third edition. Multiple sclerosis. 2020;26(14):1816–21.

4. Versini M, Jeandel PY, Rosenthal E, Shoenfeld Y. Obesity in autoimmune diseases: not a passive bystander. Autoimmun Rev. 2014;13(9):981–1000.

5. Ivancovsky-Wajcman D, Fliss-Isakov N, Webb M, Bentov I, Shibolet O, Kariv R, et al. Ultra-processed food is associated with features of metabolic syndrome and non-alcoholic fatty liver disease. Liver Int. 2021;41(11):2635–45.

6. Martinez Steele E, Juul F, Neri D, Rauber F, Monteiro CA. Dietary share of ultra-processed foods and metabolic syndrome in the US adult population. Prev Med. 2019;125:40–8.

7. Tavares LF, Fonseca SC, Garcia Rosa ML, Yokoo EM. Relationship between ultra-processed foods and metabolic syndrome in adolescents from a Brazilian Family Doctor Program. Public Health Nutr. 2012;15(1):82–7.

8. Fiolet T, Srour B, Sellem L, Kesse-Guyot E, Alles B, Mejean C, et al. Consumption of ultra-processed foods and cancer risk: results from NutriNet-Sante prospective cohort. BMJ. 2018;360:k322.

9. Mendonca RD, Lopes AC, Pimenta AM, Gea A, Martinez-Gonzalez MA, Bes-Rastrollo M. Ultra-Processed Food Consumption and the Incidence of Hypertension in a Mediterranean Cohort: The Seguimiento Universidad de Navarra Project. Am J Hypertens. 2017;30(4):358–66.

10. Schnabel L, Kesse-Guyot E, Alles B, Touvier M, Srour B, Hercberg S, et al. Association Between Ultraprocessed Food Consumption and Risk of Mortality Among Middle-aged Adults in France. JAMA Intern Med. 2019;179(4):490–8.

11. Hall KD, Ayuketah A, Brychta R, Cai H, Cassimatis T, Chen KY, et al. Ultra-Processed Diets Cause Excess Calorie Intake and Weight Gain: An Inpatient Randomized Controlled Trial of Ad Libitum Food Intake. Cell metabolism. 2019;30(1):67–77 e3.

12. Thorburn AN, Macia L, Mackay CR. Diet, metabolites, and “western-lifestyle” inflammatory diseases. Immunity. 2014;40(6):833–42.

13. Wahls TL, Chenard CA, Snetselaar LG. Review of Two Popular Eating Plans within the Multiple Sclerosis Community: Low Saturated Fat and Modified Paleolithic. Nutrients. 2019;11(2).

14. Stoiloudis P, Kesidou E, Bakirtzis C, Sintila SA, Konstantinidou N, Boziki M, et al. The Role of Diet and Interventions on Multiple Sclerosis: A Review. Nutrients. 2022;14(6).

15. Sato W, Yamamura T. Multiple sclerosis: Possibility of a gut environment-induced disease. Neurochemistry international. 2019;130:104475.

16. Oliveira SR, Kallaur AP, Lopes J, Colado Simao AN, Reiche EM, de Almeida ERD, et al. Insulin resistance, atherogenicity, and iron metabolism in multiple sclerosis with and without depression: Associations with inflammatory and oxidative stress biomarkers and uric acid. Psychiatry Res. 2017;250:113–20.

17. Oliveira SR, Simao AN, Kallaur AP, de Almeida ER, Morimoto HK, Lopes J, et al. Disability in patients with multiple sclerosis: influence of insulin resistance, adiposity, and oxidative stress. Nutrition. 2014;30(3):268–73.

18. Glatigny S, Bettelli E. Experimental Autoimmune Encephalomyelitis (EAE) as Animal Models of Multiple Sclerosis (MS). Cold Spring Harb Perspect Med. 2018;8(11).

19. Brownlow ML, Benner L, D’Agostino D, Gordon MN, Morgan D. Ketogenic diet improves motor performance but not cognition in two mouse models of Alzheimer’s pathology. PLoS One. 2013;8(9):e75713.

20. Piccio L, Stark JL, Cross AH. Chronic calorie restriction attenuates experimental autoimmune encephalomyelitis. J Leukoc Biol. 2008;84(4):940–8.

21. Bai M, Wang Y, Han R, Xu L, Huang M, Zhao J, et al. Intermittent caloric restriction with a modified fasting-mimicking diet ameliorates autoimmunity and promotes recovery in a mouse model of multiple sclerosis. J Nutr Biochem. 2021;87:108493.

22. Choi IY, Piccio L, Childress P, Bollman B, Ghosh A, Brandhorst S, et al. A Diet Mimicking Fasting Promotes Regeneration and Reduces Autoimmunity and Multiple Sclerosis Symptoms. Cell Rep. 2016;15(10):2136–46.

23. Kafami L, Raza M, Razavi A, Mirshafiey A, Movahedian M, Khorramizadeh MR. Intermittent feeding attenuates clinical course of experimental autoimmune encephalomyelitis in C57BL/6 mice. Avicenna J Med Biotechnol. 2010;2(1):47–52.

24. Razeghi Jahromi S, Ghaemi A, Alizadeh A, Sabetghadam F, Moradi Tabriz H, Togha M. Effects of Intermittent Fasting on Experimental Autoimune Encephalomyelitis in C57BL/6 Mice. Iran J Allergy Asthma Immunol. 2016;15(3):212–9.

25. Duking T, Spieth L, Berghoff SA, Piepkorn L, Schmidke AM, Mitkovski M, et al. Ketogenic diet uncovers differential metabolic plasticity of brain cells. Sci Adv. 2022;8(37):eabo7639.

26. Kim DY, Hao J, Liu R, Turner G, Shi FD, Rho JM. Inflammation-mediated memory dysfunction and effects of a ketogenic diet in a murine model of multiple sclerosis. PLoS One. 2012;7(5):e35476.

27. Martin-McGill KJ, Jackson CF, Bresnahan R, Levy RG, Cooper PN. Ketogenic diets for drug-resistant epilepsy. Cochrane Database Syst Rev. 2018;11:CD001903.

28. Wheless JW. History of the ketogenic diet. Epilepsia. 2008;49 Suppl 8:3–5.

29. Abbasi J. Interest in the Ketogenic Diet Grows for Weight Loss and Type 2 Diabetes. JAMA. 2018;319(3):215–7.

30. Athinarayanan SJ, Adams RN, Hallberg SJ, McKenzie AL, Bhanpuri NH, Campbell WW, et al. Long-Term Effects of a Novel Continuous Remote Care Intervention Including Nutritional Ketosis for the Management of Type 2 Diabetes: A 2-Year Non-randomized Clinical Trial. Front Endocrinol (Lausanne). 2019;10:348.

31. Saslow LR, Daubenmier JJ, Moskowitz JT, Kim S, Murphy EJ, Phinney SD, et al. Twelve-month outcomes of a randomized trial of a moderate-carbohydrate versus very low-carbohydrate diet in overweight adults with type 2 diabetes mellitus or prediabetes. Nutr Diabetes. 2017;7(12):304.

32. Storoni M, Plant GT. The Therapeutic Potential of the Ketogenic Diet in Treating Progressive Multiple Sclerosis. Mult Scler Int. 2015;2015:681289.

33. Wilhelm C, Surendar J, Karagiannis F. Enemy or ally? Fasting as an essential regulator of immune responses. Trends in immunology. 2021;42(5):389–400.

34. Bahr LS, Bock M, Liebscher D, Bellmann-Strobl J, Franz L, Pruss A, et al. Ketogenic diet and fasting diet as Nutritional Approaches in Multiple Sclerosis (NAMS): protocol of a randomized controlled study. Trials. 2020;21(1):3.

35. Larabee CM, Desai S, Agasing A, Georgescu C, Wren JD, Axtell RC, et al. Loss of Nrf2 exacerbates the visual deficits and optic neuritis elicited by experimental autoimmune encephalomyelitis. Mol Vis. 2016;22:1503–13.

36. Larabee CM, Hu Y, Desai S, Georgescu C, Wren JD, Axtell RC, et al. Myelin-specific Th17 cells induce severe relapsing optic neuritis with irreversible loss of retinal ganglion cells in C57BL/6 mice. Mol Vis. 2016;22:332–41.

37. Zyla K, Larabee CM, Georgescu C, Berkley C, Reyna T, Plafker SM. Dimethyl fumarate mitigates optic neuritis. Mol Vis. 2019;25:446–61.

38. Bligh EG, Dyer WJ. A rapid method of total lipid extraction and purification. Can J Biochem Physiol. 1959;37(8):911–7.

39. Li F, Marchette LD, Brush RS, Elliott MH, Le YZ, Henry KA, et al. DHA does not protect ELOVL4 transgenic mice from retinal degeneration. Mol Vis. 2009;15:1185–93.

40. Agbaga MP, Merriman DK, Brush RS, Lydic TA, Conley SM, Naash MI, et al. Differential composition of DHA and very-long-chain PUFAs in rod and cone photoreceptors. J Lipid Res. 2018;59(9):1586–96.

41. Yu M, Benham A, Logan S, Brush RS, Mandal MNA, Anderson RE, et al. ELOVL4 protein preferentially elongates 20:5n3 to very long chain PUFAs over 20:4n6 and 22:6n3. J Lipid Res. 2012;53(3):494–504.

42. Axtell RC, de Jong BA, Boniface K, van der Voort LF, Bhat R, De Sarno P, et al. T helper type 1 and 17 cells determine efficacy of interferon-beta in multiple sclerosis and experimental encephalomyelitis. Nat Med. 2010;16(4):406–12.

43. Laird NM, Ware JH. Random-effects models for longitudinal data. Biometrics. 1982;38(4):963–74.

44. Lindstrom ML, Bates DM. Nonlinear mixed effects models for repeated measures data. Biometrics. 1990;46(3):673–87.

45. Goldberg EL, Asher JL, Molony RD, Shaw AC, Zeiss CJ, Wang C, et al. beta-Hydroxybutyrate Deactivates Neutrophil NLRP3 Inflammasome to Relieve Gout Flares. Cell Rep. 2017;18(9):2077–87.

46. Salberg S, Weerwardhena H, Collins R, Reimer RA, Mychasiuk R. The behavioural and pathophysiological effects of the ketogenic diet on mild traumatic brain injury in adolescent rats. Behav Brain Res. 2019;376:112225.

47. Haghikia A, Jorg S, Duscha A, Berg J, Manzel A, Waschbisch A, et al. Dietary Fatty Acids Directly Impact Central Nervous System Autoimmunity via the Small Intestine. Immunity. 2015;43(4):817–29.

48. Sanna V, Di Giacomo A, La Cava A, Lechler RI, Fontana S, Zappacosta S, et al. Leptin surge precedes onset of autoimmune encephalomyelitis and correlates with development of pathogenic T cell responses. J Clin Invest. 2003;111(2):241–50.

49. Miller VJ, Villamena FA, Volek JS. Nutritional Ketosis and Mitohormesis: Potential Implications for Mitochondrial Function and Human Health. J Nutr Metab. 2018;2018:5157645.

50. Kale N. Optic neuritis as an early sign of multiple sclerosis. Eye Brain. 2016;8:195–202.

51. Papp V, Magyari M, Aktas O, Berger T, Broadley SA, Cabre P, et al. Worldwide Incidence and Prevalence of Neuromyelitis Optica: A Systematic Review. Neurology. 2021;96(2):59–77.

52. Reindl M, Rostasy K. MOG antibody-associated diseases. Neurology(R) neuroimmunology & neuroinflammation. 2015;2(1):e60.

53. The clinical profile of optic neuritis. Experience of the Optic Neuritis Treatment Trial. Optic Neuritis Study Group. Arch Ophthalmol. 1991;109(12):1673–8.

54. Atkins EJ, Biousse V, Newman NJ. The natural history of optic neuritis. Rev Neurol Dis. 2006;3(2):45–56.

55. Guo J, Li B, Wang J, Guo R, Tian Y, Song S, et al. Protective effect and mechanism of nicotinamide adenine dinucleotide against optic neuritis in mice with experimental autoimmune encephalomyelitis. Int Immunopharmacol. 2021;98:107846.

56. Hobom M, Storch MK, Weissert R, Maier K, Radhakrishnan A, Kramer B, et al. Mechanisms and time course of neuronal degeneration in experimental autoimmune encephalomyelitis. Brain Pathol. 2004;14(2):148–57.

57. Sattler MB, Merkler D, Maier K, Stadelmann C, Ehrenreich H, Bahr M, et al. Neuroprotective effects and intracellular signaling pathways of erythropoietin in a rat model of multiple sclerosis. Cell Death Differ. 2004;11 Suppl 2:S181–92.

58. Shindler KS, Guan Y, Ventura E, Bennett J, Rostami A. Retinal ganglion cell loss induced by acute optic neuritis in a relapsing model of multiple sclerosis. Multiple sclerosis. 2006;12(5):526–32.

59. Shindler KS, Ventura E, Dutt M, Rostami A. Inflammatory demyelination induces axonal injury and retinal ganglion cell apoptosis in experimental optic neuritis. Exp Eye Res. 2008;87(3):208–13.

60. Green AJ, McQuaid S, Hauser SL, Allen IV, Lyness R. Ocular pathology in multiple sclerosis: retinal atrophy and inflammation irrespective of disease duration. Brain. 2010;133(Pt 6):1591–601.

61. Zahoor I, Giri S. Specialized Pro-Resolving Lipid Mediators: Emerging Therapeutic Candidates for Multiple Sclerosis. Clinical reviews in allergy & immunology. 2021;60(2):147–63.

62. Hammarstrom S. Conversion of 5,8,11-eicosatrienoic acid to leukotrienes C3 and D3. J Biol Chem. 1981;256(5):2275–9.

63. Hulbert AJ, Kelly MA, Abbott SK. Polyunsaturated fats, membrane lipids and animal longevity. J Comp Physiol B. 2014;184(2):149–66.

64. Burdge GC, Wootton SA. Conversion of alpha-linolenic acid to eicosapentaenoic, docosapentaenoic and docosahexaenoic acids in young women. Br J Nutr. 2002;88(4):411–20.

65. El-behi M, Rostami A, Ciric B. Current views on the roles of Th1 and Th17 cells in experimental autoimmune encephalomyelitis. J Neuroimmune Pharmacol. 2010;5(2):189–97.

66. Imler TJ, Jr., Petro TM. Decreased severity of experimental autoimmune encephalomyelitis during resveratrol administration is associated with increased IL-17+IL-10+ T cells, CD4(-) IFN-gamma+ cells, and decreased macrophage IL-6 expression. Int Immunopharmacol. 2009;9(1):134–43.

67. Garbow JR, Doherty JM, Schugar RC, Travers S, Weber ML, Wentz AE, et al. Hepatic steatosis, inflammation, and ER stress in mice maintained long term on a very low-carbohydrate ketogenic diet. Am J Physiol Gastrointest Liver Physiol. 2011;300(6):G956–67.

68. Jornayvaz FR, Jurczak MJ, Lee HY, Birkenfeld AL, Frederick DW, Zhang D, et al. A high-fat, ketogenic diet causes hepatic insulin resistance in mice, despite increasing energy expenditure and preventing weight gain. Am J Physiol Endocrinol Metab. 2010;299(5):E808–15.

69. Zhang D, Jin W, Wu R, Li J, Park SA, Tu E, et al. High Glucose Intake Exacerbates Autoimmunity through Reactive-Oxygen-Species-Mediated TGF-beta Cytokine Activation. Immunity. 2019.

70. Westman EC, Yancy WS, Jr., Mavropoulos JC, Marquart M, McDuffie JR. The effect of a low-carbohydrate, ketogenic diet versus a low-glycemic index diet on glycemic control in type 2 diabetes mellitus. Nutr Metab (Lond). 2008;5:36.

71. Forsythe CE, Phinney SD, Fernandez ML, Quann EE, Wood RJ, Bibus DM, et al. Comparison of low fat and low carbohydrate diets on circulating fatty acid composition and markers of inflammation. Lipids. 2008;43(1):65–77.

72. Hallberg SJ, McKenzie AL, Williams PT, Bhanpuri NH, Peters AL, Campbell WW, et al. Effectiveness and Safety of a Novel Care Model for the Management of Type 2 Diabetes at 1 Year: An Open-Label, Non-Randomized, Controlled Study. Diabetes Ther. 2018;9(2):583–612.

73. Hyde PN, Sapper TN, Crabtree CD, LaFountain RA, Bowling ML, Buga A, et al. Dietary carbohydrate restriction improves metabolic syndrome independent of weight loss. JCI Insight. 2019;4(12).

74. Volek JS, Phinney SD, Krauss RM, Johnson RJ, Saslow LR, Gower B, et al. Alternative Dietary Patterns for Americans: Low-Carbohydrate Diets. Nutrients. 2021;13(10).

75. Bisht B, Darling WG, Grossmann RE, Shivapour ET, Lutgendorf SK, Snetselaar LG, et al. A multimodal intervention for patients with secondary progressive multiple sclerosis: feasibility and effect on fatigue. J Altern Complement Med. 2014;20(5):347–55.

76. Bisht B, Darling WG, Shivapour ET, Lutgendorf SK, Snetselaar LG, Chenard CA, et al. Multimodal intervention improves fatigue and quality of life in subjects with progressive multiple sclerosis: a pilot study. Degener Neurol Neuromuscul Dis. 2015;5:19–35.

77. Bisht B, Darling WG, White EC, White KA, Shivapour ET, Zimmerman MB, et al. Effects of a multimodal intervention on gait and balance of subjects with progressive multiple sclerosis: a prospective longitudinal pilot study. Degener Neurol Neuromuscul Dis. 2017;7:79–93.

78. Fellows Maxwell K, Wahls T, Browne RW, Rubenstein L, Bisht B, Chenard CA, et al. Lipid profile is associated with decreased fatigue in individuals with progressive multiple sclerosis following a diet-based intervention: Results from a pilot study. PLoS One. 2019;14(6):e0218075.

79. Irish AK, Erickson CM, Wahls TL, Snetselaar LG, Darling WG. Randomized control trial evaluation of a modified Paleolithic dietary intervention in the treatment of relapsing-remitting multiple sclerosis: a pilot study. Degener Neurol Neuromuscul Dis. 2017;7:1–18.

80. Lee JE, Bisht B, Hall MJ, Rubenstein LM, Louison R, Klein DT, et al. A Multimodal, Nonpharmacologic Intervention Improves Mood and Cognitive Function in People with Multiple Sclerosis. J Am Coll Nutr. 2017;36(3):150–68.

81. Wahls TL, Titcomb TJ, Bisht B, Eyck PT, Rubenstein LM, Carr LJ, et al. Impact of the Swank and Wahls elimination dietary interventions on fatigue and quality of life in relapsing-remitting multiple sclerosis: The WAVES randomized parallel-arm clinical trial. Mult Scler J Exp Transl Clin. 2021;7(3):20552173211035399.

82. Bjornevik K, Chitnis T, Ascherio A, Munger KL. Polyunsaturated fatty acids and the risk of multiple sclerosis. Multiple sclerosis. 2017;23(14):1830–8.

83. Penkert H, Bertrand A, Tiwari V, Breimann S, Muller SA, Jordan PM, et al. Proteomic and lipidomic profiling of demyelinating lesions identifies fatty acids as modulators in lesion recovery. Cell Rep. 2021;37(4):109898.

84. Poisson LM, Suhail H, Singh J, Datta I, Denic A, Labuzek K, et al. Untargeted Plasma Metabolomics Identifies Endogenous Metabolite with Drug-like Properties in Chronic Animal Model of Multiple Sclerosis. J Biol Chem. 2015;290(52):30697–712.

85. Kooij G, Troletti CD, Leuti A, Norris PC, Riley I, Albanese M, et al. Specialized pro-resolving lipid mediators are differentially altered in peripheral blood of patients with multiple sclerosis and attenuate monocyte and blood-brain barrier dysfunction. Haematologica. 2020;105(8):2056–70.

86. Villoslada P, Alonso C, Agirrezabal I, Kotelnikova E, Zubizarreta I, Pulido-Valdeolivas I, et al. Metabolomic signatures associated with disease severity in multiple sclerosis. Neurology(R) neuroimmunology & neuroinflammation. 2017;4(2):e321.

87. Rumble JM, Huber AK, Krishnamoorthy G, Srinivasan A, Giles DA, Zhang X, et al. Neutrophil-related factors as biomarkers in EAE and MS. J Exp Med. 2015;212(1):23–35.

88. Aube B, Levesque SA, Pare A, Chamma E, Kebir H, Gorina R, et al. Neutrophils mediate blood-spinal cord barrier disruption in demyelinating neuroinflammatory diseases. J Immunol. 2014;193(5):2438–54.

89. Jiang W, St-Pierre S, Roy P, Morley BJ, Hao J, Simard AR. Infiltration of CCR2+Ly6Chigh Proinflammatory Monocytes and Neutrophils into the Central Nervous System Is Modulated by Nicotinic Acetylcholine Receptors in a Model of Multiple Sclerosis. J Immunol. 2016;196(5):2095–108.

90. Levesque SA, Pare A, Mailhot B, Bellver-Landete V, Kebir H, Lecuyer MA, et al. Myeloid cell transmigration across the CNS vasculature triggers IL-1beta-driven neuroinflammation during autoimmune encephalomyelitis in mice. J Exp Med. 2016;213(6):929–49.

91. Steinbach K, Piedavent M, Bauer S, Neumann JT, Friese MA. Neutrophils amplify autoimmune central nervous system infiltrates by maturing local APCs. J Immunol. 2013;191(9):4531–9.

92. Yan Z, Yang W, Parkitny L, Gibson SA, Lee KS, Collins F, et al. Deficiency of Socs3 leads to brain-targeted EAE via enhanced neutrophil activation and ROS production. JCI Insight. 2019;5.

93. Ghafouri-Fard S, Honarmand K, Taheri M. A comprehensive review on the role of chemokines in the pathogenesis of multiple sclerosis. Metab Brain Dis. 2021;36(3):375–406.

94. Tsutsui M, Hirase R, Miyamura S, Nagayasu K, Nakagawa T, Mori Y, et al. TRPM2 Exacerbates Central Nervous System Inflammation in Experimental Autoimmune Encephalomyelitis by Increasing Production of CXCL2 Chemokines. J Neurosci. 2018;38(39):8484–95.

95. Carlson T, Kroenke M, Rao P, Lane TE, Segal B. The Th17-ELR+ CXC chemokine pathway is essential for the development of central nervous system autoimmune disease. J Exp Med. 2008;205(4):811–23.

96. Matejuk A, Dwyer J, Ito A, Bruender Z, Vandenbark AA, Offner H. Effects of cytokine deficiency on chemokine expression in CNS of mice with EAE. J Neurosci Res. 2002;67(5):680–8.

97. Stoolman JS, Duncker PC, Huber AK, Giles DA, Washnock-Schmid JM, Soulika AM, et al. An IFNgamma/CXCL2 regulatory pathway determines lesion localization during EAE. Journal of neuroinflammation. 2018;15(1):208.

98. Huang J, Khademi M, Fugger L, Lindhe O, Novakova L, Axelsson M, et al. Inflammation-related plasma and CSF biomarkers for multiple sclerosis. Proc Natl Acad Sci U S A. 2020;117(23):12952–60.

99. Adzemovic MZ, Ockinger J, Zeitelhofer M, Hochmeister S, Beyeen AD, Paulson A, et al. Expression of Ccl11 associates with immune response modulation and protection against neuroinflammation in rats. PLoS One. 2012;7(7):e39794.

100. Ruppova K, Lim JH, Fodelianaki G, August A, Neuwirth A. Eosinophils are dispensable for development of MOG(35-55)-induced experimental autoimmune encephalomyelitis in mice. Immunol Lett. 2021;239:72–6.

101. Nazarinia D, Behzadifard M, Gholampour J, Karimi R, Gholampour M. Eotaxin-1 (CCL11) in neuroinflammatory disorders and possible role in COVID-19 neurologic complications. Acta Neurol Belg. 2022;122(4):865–9.

102. Erta M, Quintana A, Hidalgo J. Interleukin-6, a major cytokine in the central nervous system. Int J Biol Sci. 2012;8(9):1254–66.

103. Maimone D, Guazzi GC, Annunziata P. IL-6 detection in multiple sclerosis brain. J Neurol Sci. 1997;146(1):59–65.

104. Stampanoni Bassi M, Iezzi E, Drulovic J, Pekmezovic T, Gilio L, Furlan R, et al. IL-6 in the Cerebrospinal Fluid Signals Disease Activity in Multiple Sclerosis. Frontiers in cellular neuroscience. 2020;14:120.

105. Setiadi AF, Abbas AR, Jeet S, Wong K, Bischof A, Peng I, et al. IL-17A is associated with the breakdown of the blood-brain barrier in relapsing-remitting multiple sclerosis. Journal of neuroimmunology. 2019;332:147–54.

106. Bettelli E, Carrier Y, Gao W, Korn T, Strom TB, Oukka M, et al. Reciprocal developmental pathways for the generation of pathogenic effector TH17 and regulatory T cells. Nature. 2006;441(7090):235–8.

107. Mangan PR, Harrington LE, O’Quinn DB, Helms WS, Bullard DC, Elson CO, et al. Transforming growth factor-beta induces development of the T(H)17 lineage. Nature. 2006;441(7090):231–4.

108. Serada S, Fujimoto M, Mihara M, Koike N, Ohsugi Y, Nomura S, et al. IL-6 blockade inhibits the induction of myelin antigen-specific Th17 cells and Th1 cells in experimental autoimmune encephalomyelitis. Proc Natl Acad Sci U S A. 2008;105(26):9041–6.

109. Veldhoen M, Hocking RJ, Atkins CJ, Locksley RM, Stockinger B. TGFbeta in the context of an inflammatory cytokine milieu supports de novo differentiation of IL-17-producing T cells. Immunity. 2006;24(2):179–89.

110. Ma S, Huang Q, Tominaga T, Liu C, Suzuki K. An 8-Week Ketogenic Diet Alternated Interleukin-6, Ketolytic and Lipolytic Gene Expression, and Enhanced Exercise Capacity in Mice. Nutrients. 2018;10(11).

111. Nakamura K, Tonouchi H, Sasayama A, Ashida K. A Ketogenic Formula Prevents Tumor Progression and Cancer Cachexia by Attenuating Systemic Inflammation in Colon 26 Tumor-Bearing Mice. Nutrients. 2018;10(2).

112. Nandivada P, Fell GL, Pan AH, Nose V, Ling PR, Bistrian BR, et al. Eucaloric Ketogenic Diet Reduces Hypoglycemia and Inflammation in Mice with Endotoxemia. Lipids. 2016;51(6):703–14.

113. Norwitz NG, Winwood R, Stubbs BJ, D’Agostino DP, Barnes PJ. Case Report: Ketogenic Diet Is Associated With Improvements in Chronic Obstructive Pulmonary Disease. Front Med (Lausanne). 2021;8:699427.

114. Thambi M, Nathan J, Bailur S, Unnikrishnan MK, Ballal M, Radhakrishnan K. Is the antiseizure effect of ketogenic diet in children with drug-resistant epilepsy mediated through proinflammatory cytokines? Epilepsy Res. 2021;176:106724.

115. Yang X, Cheng B. Neuroprotective and anti-inflammatory activities of ketogenic diet on MPTP-induced neurotoxicity. Journal of molecular neuroscience: MN. 2010;42(2):145–53.

116. Zhu Y, Tang X, Cheng Z, Dong Q, Ruan G. The Anti-Inflammatory Effect of Preventive Intervention with Ketogenic Diet Mediated by the Histone Acetylation of mGluR5 Promotor Region in Rat Parkinson’s Disease Model: A Dual-Tracer PET Study. Parkinson’s disease. 2022;2022:3506213.

117. Bertoli S, Neri IG, Trentani C, Ferraris C, De Amicis R, Battezzati A, et al. Short-term effects of ketogenic diet on anthropometric parameters, body fat distribution, and inflammatory cytokine production in GLUT1 deficiency syndrome. Nutrition. 2015;31(7-8):981–7.

118. Fraser DA, Thoen J, Djoseland O, Forre O, Kjeldsen-Kragh J. Serum levels of interleukin-6 and dehydroepiandrosterone sulphate in response to either fasting or a ketogenic diet in rheumatoid arthritis patients. Clin Exp Rheumatol. 2000;18(3):357–62.

119. Vidali S, Aminzadeh-Gohari S, Feichtinger RG, Vatrinet R, Koller A, Locker F, et al. The ketogenic diet is not feasible as a therapy in a CD-1 nu/nu mouse model of renal cell carcinoma with features of Stauffer’s syndrome. Oncotarget. 2017;8(34):57201–15.

120. Longo VD, Panda S. Fasting, Circadian Rhythms, and Time-Restricted Feeding in Healthy Lifespan. Cell metabolism. 2016;23(6):1048–59.

121. Luis D, Zlatkis K, Comenge B, Garcia Z, Navarro JF, Lorenzo V, et al. Dietary Quality and Adherence to Dietary Recommendations in Patients Undergoing Hemodialysis. J Ren Nutr. 2016;26(3):190–5.

122. Mellor R, Saunders-Dow E, Mayr HL. Scope of Use and Effectiveness of Dietary Interventions for Improving Health-Related Outcomes in Veterans: A Systematic Review. Nutrients. 2022;14(10).

