## Supplemental Figure legends for "Dietary Protection Against the Visual and Motor Deficits Induced by Experimental Autoimmune Encephalomyelitis"

**Supplemental Figure 1.** **Diet compositions and weight changes. (A)** Complete ingredients of the custom-formulated purified KD and CD diets. **(B)** Repeated measures graph showing patterns of weight changes in mice fed the KD or CD in the absence of EAE. For each individual mouse (n=5/diet), weights are graphed at baseline (circles) and 5 weeks post-feeding of the indicated diet (squares). (**C**) Repeated measures graphs comparing baseline and final weights of female (left graph) and male (right graph) mice consuming a KD or CD during EAE. Weight comparisons span from the day of immunization (0 dpi; circles) to the termination of experiment (21 dpi; squares). **(D)** Shown are tabulated average glucose and ketone readings taken at baseline, 0 dpi, and 21 dpi, along with standard deviations (SD) and p values. Statistical comparisons of KD versus CD for females (F) and males (M) done by unpaired T test. For (B-D), * p<0.05, ** p<0.01, **** p<0.0001.)

**Supplemental Figure 2. The KD preserves vision and RGCs in female and male mice. (A)** OKT measurements of visual acuity of the more affected (MA; top graph) and less affected (LA; bottom graph) eyes in female mice on the indicated diets. n=14-17 mice/diet. **(B)** Same as (A) for male mice. n=10-12 mice/diet. For graphs in (**A**) and (**B**), whisker-bar standard deviations and p-values for differences between curves were computed with mixed linear model implementation of lme function, nlme R package. For LA graphs, insets with adjusted scales of x-axis and y-axis highlight significant differences in visual acuity between diets. (**C**) Graph of Brn3A**^+^** RGC cell counts in central, medial, and peripheral retinas of MA eyes as a function of sex and diet. 2-way ANNOVA was performed to determine statistical significance. ‘ns’ denotes not significant. * p<0.05, ** p<0.01, *** p<0.001, **** p<0.0001. Data compiled $\geq$ 4 independent experiments.

**Supplemental Figure 3. The KD preserves optic nerve myelination and reduces infiltrates.** Mean values and 95% confidence intervals for CNPase (**A**), Iba1 (**B**), and CD3 (**C**) labeling of optic nerve sections. Data shown for females and males separately (‘F’; ‘M’) as well as combined (‘F+M’). Whisker bars depict 95% confidence intervals for each variable and horizontal bars with accompanying p-values above the bars indicate statistical significance.

**Supplemental Figure 4. Fatty acids analyzed in the plasma and diets. (A,B)** Graphs depicting nmols of individual fatty acids per mg of protein in the plasma that are statistically significantly different (*p$\leq$0.05, **p<0.01, ***p<0.001, ****p<0.0001 as determined by Mann-Whitney test with Bonferroni correction) between EAE mice fed the KD versus the CD. Graphs represent data from females (**A**) and males (**B**). **(C)** Graph showing micrograms ($\mu$g) of each fatty acid per milligram (mg) of diet comparing the KD, CD, and standard chow (SC). For each graph, the fatty acid omega designations are listed on the x-axis. n$\geq$6 mice per diet from $\geq$2 independent experiments and 3 diet samples.

**Supplemental Figure 5. Cytokines analyzed in the plasma.** Graphs showing concentrations of the indicated cytokines and chemokines in the plasma 21 dpi as a function of both sex and diet. Female and male EAE mice consumed either the KD (blue for females and green for males) or a CD (red for females and purple for males), 4-8 mice per diet per sex from $\geq$ 2 independent experiments. Statistical significance was determined by Mann-Whitney test with ‘ns’ indicating not significant. * p<0.05, ** p<0.01, *** p<0.001, **** p<0.0001.

**Supplemental Figure 6. EAE mice consuming a KD mount a T cell response to MOG antigen immunization.**  **(A)** Graphs of % live splenocytes positive for the indicated markers above each graph following *ex vivo* stimulation with MOG_35-55_ peptide for 24 hrs, with addition of monensin, PMA, and ionomycin for the final 5 hrs. Unstimulated cells are shown for comparison. n=13-20; p-values calculated by unpaired parametric t-test; ns = not significant; **p$\leq$0.01. **(B)** Representative examples of gating strategy on live CD3^+^CD4^+^CD44^++^ cells. Expression of IL-17 and IFNγ was assessed in CD3^+^CD4^+^CD44^++^ splenocytes by flow cytometry. **(C)** Examples of gating strategy for live, CD3^+^CD4^+^ CD44^++^ T cells showing similar levels of activated splenic T cells for mice fed both diets.

**Supplemental Figure 7. KD fed mice have decreased incidence of EAE and disease severity. (A)** Table showing the incidence of “breakthrough” mice on the KD that developed EAE motor and visual deficits within 2 days of harvest (i.e., 19-21 dpi). Data compiled $\geq$ 4 independent experiments. **(B)** Kaplan Meier graph showing the percent of mice (females + males) that developed EAE symptoms as a function of diet and time post-immunization. Mice on the KD were followed for 35 days after immunization. N=18 mice on KD (green trace) and n=10 mice on CD (orange trace). Statistical significance determined by log-rank (Mantel-Cox) test; ****p<0.0001. **(C)** Graph of mean EAE motor scores as a function of diet using symptomatic mice from (**B**). N=5 mice on KD and n=8 mice on CD that manifested symptoms. Statistical significance determined by unpaired T test; ***p<0.001.
